# A gradient of electrophysiological novelty responses along the human hippocampal long axis

**DOI:** 10.1101/2021.11.02.466832

**Authors:** Mar Yebra, Ole Jensen, Lukas Kunz, Stephan Moratti, Nikolai Axmacher, Bryan A Strange

**Author notes:** equal contribution (Nikolai Axmacher, Bryan A Strange). Ueli Rutishauser Lab. Department of Neurosurgery, Cedars-Sinai Medical Center, Los Angeles, California 90048, US.

## Abstract

The hippocampus is implicated in novelty detection, thought to be important for regulating entry of information into long-term memory. Whether electrophysiological responses to novelty differ along the human hippocampal long axis is currently unknown. By recording from electrodes implanted longitudinally in the hippocampus of epilepsy patients, here we show a gradual increase of theta frequency oscillatory power from anterior to posterior in response to unexpected stimuli, superimposed on novelty responses common to all long axis portions. Intracranial event-related potentials (iERPs) were larger for unexpected *vs*. expected stimuli and demonstrated a polarity inversion between the hippocampal head (HH) and body (HB). We observed stronger theta coherence between HH and hippocampal tail (HT) than between HB and HT, similarly for expected and unexpected stimuli. This was accompanied by theta and alpha traveling waves with surprisingly variable direction of travel characterized by a ∼180° phase lag between hippocampal poles. Interestingly, this phase lag showed a pronounced phase offset between anterior and middle (HH-HB) hippocampal portions coinciding anatomically with a drop in theta coherence and the novelty iERP polarity inversion. Our findings indicate common response properties along the hippocampal long axis to unexpected stimuli, as well as a multifaceted, non-uniform engagement along the long axis for novelty processing.

## Introduction

Distinct patterns of functional organization along the hippocampal long-axis are evident in rodents and primates, including humans, providing a potential framework for understanding the multiple functions ascribed to the hippocampus. Functional differences along its long axis were apparent from early rodent studies (Moser & Moser, 1998; Nadel, 1968) and are increasingly studied in human participants (Brunec et al., 2018; Collin, Milivojevic, & Doeller, 2017; Moser & Moser, 1998; Poppenk, Evensmoen, Moscovitch, & Nadel, 2013; Schlichting, Mumford, & Preston, 2015; Strange, Witter, Lein, & Moser, 2014; B. Strange, Fletcher, Henson, Friston, & Dolan, 1999; Zeidman & Maguire, 2016). In rodents, a dichotomous view has predominated, which implicated the ventral hippocampus (the rodent homologue of the anterior hippocampus in humans) in emotional functions, while the dorsal hippocampus (posterior hippocampus in humans) was proposed to mainly mediate spatial and memory functions (Fanselow & Dong, 2010). Intrinsic connectivity within the rodent (Amaral & Witter, 1989; Fricke & Cowan, 1978; Ishizuka, Weber, & Amaral, 1990; Li, Somogyi, Ylinen, & Buzsáki, 1994; Swanson, Wyss, & Cowan, 1978), and to a lesser extent non-human primate (Kondo, Lavenex, & Amaral, 2009) hippocampal long axis, shows an abrupt change from the ventral third to the dorsal two-thirds. By contrast, electrophysiological recordings in rodents show gradually increasing place field size from dorsal to ventral hippocampus (Kjelstrup et al., 2008), suggesting that the long axis is organised along a gradient, a pattern also evident in extrinsic anatomical connectivity (Strien, Cappaert, & Witter, 2009). Thus, an emerging view is that the hippocampal long axis shows superimposed graded and discretised levels of organisation (Strange et al., 2014).

In humans, evidence for graded *vs*. discrete functional differences along the hippocampal long axis is currently lacking. An early observation from functional MRI in humans was that responses in anterior portions of the hippocampus were greater to novel or unexpected stimuli than in posterior portions, with the anterior hippocampus showing response habituation with repeated exposure to these stimuli (Strange et al., 1999). This dissociation has been replicated in a number of human neuroimaging studies, indicating a novelty effect primarily in the anterior hippocampus (Constable et al., 2000; Dolan & Fletcher, 1997; see Grady, 2020 for review; Haxby et al., 1996; Kohler, Crane, & Milner, 2002; Saykin et al., 1999; B. Strange et al., 1999; Tulving, Markowitsch, Craik, Habib, & Houle, 1996)). Whether the anterior hippocampal sensitivity to novelty is demarcated to this specific portion, or shows a gradual change over the long axis, is currently unknown.

Novel, or unexpected stimuli or events are preferentially encoded into memory as compared to expected ones (Ranganath & Rainer, 2003; Von Restorff, 1933) and a hippocampal role in detecting mismatches between expectation and experience was described in early reports from animal models (O’keefe & Nadel, 1978; Ranck, 1973; Vinogradova, 1975). Novelty responses in humans have been widely studied using oddball paradigms, where the oddball stimuli represent violations of expectation due to their deviance from the prevailing context (Rugg, 1995). There is electrophysiological evidence for a hippocampal role in oddball detection from intracranial (Barbeau, Chauvel, Moulin, Regis, & Liegeois-Chauvel, 2017; Halgren et al., 1980; Rosburg et al., 2007; Smith et al., 1990) and scalp (Garrido, Barnes, Kumaran, Maguire, & Dolan, 2015; Knight, 1996) EEG recordings in humans. Large hippocampal intracranially-recorded event-related potentials (iERP) to unexpected *vs*. expected stimuli have been termed “MTL P300” in reference to the novelty P300 (Kutas, McCarthy, & Donchin, 1977), a positive scalp EEG component recorded in similar experimental conditions (Brázdil, Rektor, Daniel, Dufek, & Jurák, 2001; Fell et al., 2005; Halgren et al., 1995; Halgren et al., 1980; McCarthy, Wood, Williamson, & Spencer, 1989). With respect to oscillatory responses, intracranially recorded human hippocampal responses show power difference between processing of unexpected and expected items activity in the theta and high gamma (70–90 Hz) bands (Axmacher et al., 2010). There is also evidence from iEEG recordings showing increased low-frequency power (in the slow-theta range) in the human hippocampus during associative prediction violations in a mismatch detection task (Chen et al., 2013).

These previous analyses did not, however, consider the position of the electrode contacts along the human hippocampal long axis. In rodents, recordings along the longitudinal axis of the hippocampus showed that novel space increases theta and gamma coherence across the long axis of CA1 (Penley et al., 2013). Furthermore, a travelling wave phenomenon in the theta frequency band has been observed in rodents (Patel, Fujisawa, Berenyi, Royer, & Buzsaki, 2012) and in humans (Zhang & Jacobs, 2015), suggesting a gradual change in phase along the long axis of the hippocampus. Hence, a characterisation – at a detailed oscillatory level – of how electrophysiological responses to novelty change from the hippocampal head (anterior) to its body and tail (posterior) is currently lacking, which could be key to understanding hippocampal functional organization in humans.

To address this, we examined electrophysiological responses directly recorded from the hippocampus in patients with intracranial electrodes implanted for pre-surgical evaluation of epilepsy, while these patients viewed expected and unexpected stimuli (Axmacher et al., 2010). These rare patients had undergone posterior-anterior electrode implantation, yielding multiple electrodes recording sites along the hippocampal long axis. By clustering contacts along the long axis of the hippocampus in the hippocampal head (HH), hippocampal body (HB) and hippocampal tail (HT), we specifically tested for novelty responses not only in the anterior part but also in the middle and posterior portions of the hippocampus. We used novelty-evoked iERPs locked to the onset of expected and unexpected stimuli and novelty-induced time frequency representations to explore differences in these responses along the long axis. Our data show novelty-evoked iERPs polarity inversion between the HH and HB. Conversely, we observe a theta and gamma power increase during novelty processing in all three segments, theta power being greatest in the tail. A further empirical question posed here was whether a stimulus type known to engage the hippocampus (*i.e.,* unexpected stimuli) modulated the oscillatory coupling between hippocampal long-axis portions. While we do observe theta and alpha coherence differences between hippocampal portions, these measures were unaffected by processing of information related to expectancy and were subsequently found to reflect a traveling wave phenomenon.

## Methods

### Participants

Eleven pharmacoresistant temporal lobe epilepsy patients (eight females; mean age ± SD: 37.7 ± 11.54 years) participated in this study (**Supplementary Table 1**). Recordings were performed at the Department of Epileptology, University of Bonn, Germany. Five patients suffered from unilateral hippocampal sclerosis, one presented with loss of gray-white matter differentiation in the left temporal pole, one with lesion to the left temporal pole and four showed no visible pathology. The study was approved by the local ethics committee, and all patients gave written informed consent.

### Electrode implantation

Hippocampal depth electrodes (AD-Tech, Racine, WI, USA) were implanted using a computerized tomography-based stereotactic insertion technique (Van Roost, Solymosi, Schramm, van Oosterwyck, & Eiger, 1998) following a posterior-to-anterior insertion trajectory. Nine out the eleven patients had bilateral hippocampal electrodes. Electrodes had 10 cylindrical platinum-iridium contacts, with a center-to-center distance between adjacent contacts of 4.5 mm, and a diameter of 1.3 mm.

### Electrode contact localization

Electrode localization was performed using FSL (https://fsl.fmrib.ox.ac.uk/fsl/fslwiki/FSL); (Jenkinson, Beckmann, Behrens, Woolrich, & Smith, 2012) and PyLocator (http://pylocator.thorstenkranz.de/). First, the post-implantation MR image was co-registered with the pre-implantation MR image. The pre-implantation MR image was then normalized to MNI space, applying the normalization matrix to the post-implantation MR image in parallel. Normalized post-implantation images were inspected using PyLocator and channel locations were manually identified. We then assigned probabilistic anatomic labels to the identified MNI coordinates of each electrode channel using the anatomy toolbox (version 2.2c; http://www.fz-juelich.de/inm/inm-1/DE/Forschung/_docs/SPMAnatomyToolbox/SPMAnatomyToolbox_node.html; (Eickhoff et al., 2005)) running under SPM12 (http://www.fil.ion.ucl.ac.uk/spm/software/spm12/). Note that the pre-MRI scans for patients 1 and 8 were unavailable (**Supplementary Figure 1**) thus requiring that the localization of the MNI coordinates be performed on the post-operative MRI scan in these patients.

After exclusion of electrodes (see below), each patient had 5.9 ± 1.4 (mean ± SD) remaining hippocampal contacts. The uncal apex was used as the transition point from HH to HB (Poppenk et al., 2013). Coordinates where the fimbria-fornix was visualized posterior to the pulvinar nucleus were used as the transition from HB to HT (Malykhin et al., 2007). The hippocampal tail was defined from the end of the body to the last posterior slice (**Figure 1A, B, Supplementary Figure 1**). Eight patients had electrodes in all hippocampal portions, i.e., in Hippocampal Head (HH), Hippocampal Body (HB) and Hippocampal Tail (HT), 3 patients did not have contacts in the HT.

**Figure 1.**
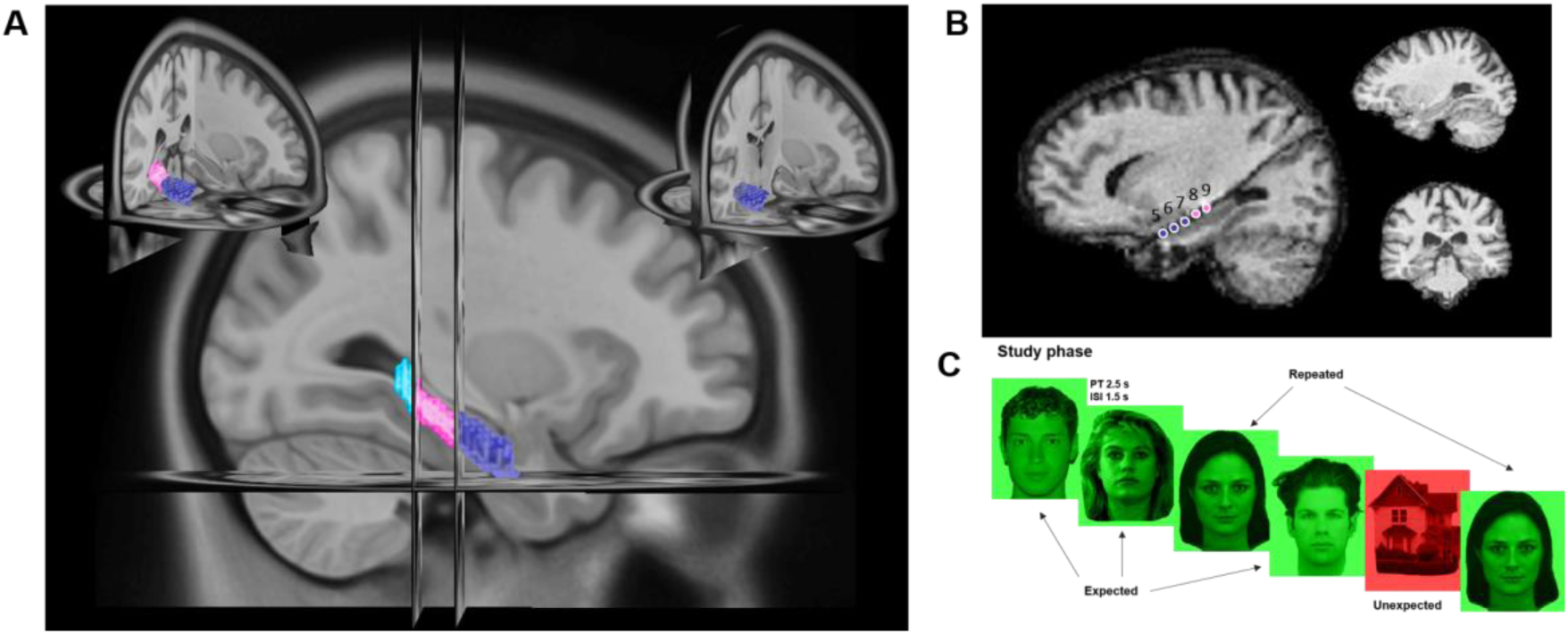
Hippocampal long-axis anatomy and experimental protocol. **A**. Segmentation criteria for hippocampal portion definition. Hippocampal Head (HH), Body (HB) and Tail (HT) are depicted in dark blue, pink and turquoise, respectively. Insets show a 3D view of the first slice of the hippocampal tail (left) and a 3D view with coronal slice at y=-21 (MNI coordinates) at the uncal apex landmark used for long-axis segmentation between HH and HB (right). **B**. Pre- and post-implantation MRI. Example of pre (two right panels) and post (left panel) MRI showing the location of the contacts along the long axis of the hippocampus for one patient. **C**. Experimental paradigm. Study Phase (encoding). The majority of presented items belonged to one category with respect to background colour and content (expected items; *e.g.*, red faces), while a minority of items were deviant (unexpected items; *e.g.*, green houses). Repeat stimuli were from the same category of expected items but had been studied during a familiarization phase and repeated throughout the experiment (PR: presentation time, ISI: Inter-Stimulus interval).

### Experimental design

The experiment was conducted across different ∼15-minute sessions including familiarization, encoding and retrieval phases (Figure 1C). During the familiarization phase, four stimuli were used as “repeated” stimuli, each of which was presented four times randomly. During encoding phases, 112 pictures from two different categories, houses or faces, were presented. 72% of the stimuli were unique “expected” items of one of the two categories, 14% of the stimuli were unique “unexpected” items from the alternative category presented as expected. 14% of the trials were “repeated” trials from the same category of expected items but they were studied during the familiarization phase and were also repeated throughout the experiment. The category of expected *vs.* unexpected items was counter-balanced over patients/sessions. During encoding, each stimulus was presented for 2500ms with a 1500ms inter-trial interval. To ensure subjects were actively participating and attending the stimuli, they were asked to rate each image as pleasant or unpleasant by pressing one of two mouse buttons with the right hand. That is, patients did not perform a typical oddball (P3b) task – making a response to rare, target deviant items – but responded to all stimuli and processed deviant items implicitly (P3a task). The order of all trials was pseudorandomized using an m-sequence (Buračas & Boynton, 2002), with the restriction that unexpected trials could never occur consecutively.

The experiment consisted of up to eight encoding sessions. Different sets of stimuli were used for each session except for the stimuli used on repeat trials. Five out of the eleven patients included in the study completed eight sessions, four of them completed four sessions, one of them five sessions and one completed three sessions. In this work only iEEG recordings from the encoding phases for the expected and unexpected conditions were analyzed. Data from this task have been reported in Axmacher et al (Axmacher et al., 2010), without analyzing the different hippocampal contacts as a function of long axis position.

### Data acquisition

Recordings were performed using a Stellate recording system (Stellate GmbH, Munich, Germany). Depth iEEG recordings were referenced to linked mastoids and recorded at a sampling rate of 1000 Hz. There was a hardware bandpass filter between 0.01 Hz (6 dB/octave) and 300 Hz (12 dB/octave).

### Data analysis

#### Pre-processing

Trials were visually inspected for artifacts (e.g., epileptiform spikes), and trials with artifacts were excluded from further analysis (number of rejected trials per patient is provided in **Supplementary Table 1**). In nine out of eleven patients, recordings were taken from the nonfocal hemisphere (i.e., contralateral to the epileptogenic focus), to minimize the possibility of artifact contamination. Two unilaterally implanted patients were included, with their recordings carefully cleaned during visual artifact rejection. After artifact rejection, contacts were labelled depending on the hippocampal portion they belonged to: HH, HB and HT, using the anatomical landmarks described above. Given that we intended to examine differences along the long axis of the hippocampus, we re-referenced our data to an average reference of all hippocampal contacts, similar to other studies (Alexander et al., 2013; Bahramisharif et al., 2013; De Munck et al., 2007; Massimini, Huber, Ferrarelli, Hill, & Tononi, 2004; Patten, Rennie, Robinson, & Gong, 2012; van der Meij, Kahana, & Maris, 2012; Zhang & Jacobs, 2015). Furthermore, this montage reportedly yields larger theta signals (Zhang & Jacobs, 2015).

#### Intracranial event-related potentials

We calculated iERPs for each patient and hippocampal contact locked to the onset of the stimuli for expected and unexpected conditions using Fieldtrip (http://www.fieldtriptoolbox.org) for Matlab (Mathworks 2014b). We defined each trial from 100 ms before the stimulus onset until 2 s after. Linear trends were removed from the data for each trial and a baseline correction (from -100 to 0 ms relative to stimulus onset) was performed. Data were filtered using a 10th order finite impulse response (FIR) low pass filter of 20 Hz. Grand averages across subjects were calculated for the different portions (HH, HB, HT) and conditions (expected, unexpected). For statistical testing, we used a cluster-based corrected (Maris & Oostenveld, 2007) two-tailed *t*-test comparing unexpected and expected conditions for each hippocampal long-axis portion 1 s after the onset of the stimuli using a maximum summation cluster statistic Montecarlo method. Specifically, we first identified individual time points with t-values corresponding to p-values smaller than 0.05, and then extracted the largest contiguous cluster by summing up t-values. The resulting summed t-value was then compared to the clusters obtained after shuffling the labels across trials (1,000 permutations). Clusters were considered significant if their summed t-values exceeded those of 975 of these surrogates, corresponding to a corrected two-tailed cluster threshold of p<0.05.

A one way ANOVA on the differences between unexpected and expected stimuli with portion (HH, HB and HT) as within-subject factor was calculated on the 8 patients who had contacts in all 3 hippocampal portions (the remaining 3 patients did not have contacts in the tail) in a time window where differences in MTL-P300 have been reported for novelty processing (250-400 ms after stimulus onset) (Axmacher et al., 2010; Chen et al., 2013; Halgren et al., 1980; Knight, 1996; Polich, 2007; Ranganath & Rainer, 2003; Smith et al., 1990; Soltani & Knight, 2000). Again, a cluster-based correction was applied as described above. Post-hoc *t*-tests were subsequently calculated between expected and unexpected conditions within each hippocampal portion and for the expected vs. unexpected difference between portions on the averaged power in the resultant significant time window.

#### Time Frequency Analysis

Condition-specific responses were analysed in the time-frequency domain using the Fieldtrip Matlab toolbox. Data were first notch-filtered (50 Hz), demeaned and detrended. In order to inspect the conditions separately, a baseline correction was applied from -100ms to -10ms relative to stimulus onset. When comparing the different conditions, no baseline correction was applied, since the relative difference between expected and unexpected condition was shown. For the lower frequencies we used a Hanning taper method with a fixed window length of 500 ms from 1 to 34 Hz in steps of 2 Hz. For the higher frequencies, from 32.5 to 150 Hz in 2.5 Hz steps, a multitaper method was used with a frequency smoothing of 10 Hz and a sliding time window of 400 ms. Trial-by-trial automatic artifact rejection was then performed in the frequency domain; the maximum of the logarithm of power per frequency bin was calculated and all trials with mean power 3 standard deviations above or below the mean were excluded from further analysis, and the same number of trials for different portions of the hippocampus were selected. Subsequently, data from contacts within each of the three different hippocampal long-axis portions (HH, HB and HT) were averaged for each patient. Portions were then grand averaged across patients. A two tailed *t*-test between unexpected and expected condition was performed for each portion. Next, a cluster-based correction (Maris & Oostenveld, 2007) was performed, using a maxsum cluster statistic Montecarlo method with a cluster alpha 0.05 and alpha of 0.025 and 1000 randomizations for lower (1-34 Hz) and higher (34-150 Hz) frequency ranges separately.

In order to find a common cluster in the difference between unexpected and expected conditions along the long-axis of the hippocampus, a one way ANOVA was calculated across the 8 patients with contacts in all 3 hippocampal portions on the difference between unexpected and expected conditions for the lower frequencies (1-20 Hz). This was performed in a time window (250-400 ms after stimulus onset) where differences in novelty processing have been previously reported in the theta and slow-theta frequency ranges (Chen et al., 2013). Again, cluster-based correction was employed. Post-hoc *t*-tests were subsequently calculated between expected and unexpected conditions within each portion and between the differences across conditions and portions on the averaged power in the resultant significant time-frequency window.

#### Coherency analysis (absolute value and phase) for theta and alpha peaks

A measure of synchrony was calculated between the contacts from the different hippocampal portions. We calculated the absolute value (Coherence) and the angle (Phase Angle) of the complex Coherency measure. To detect narrow-band oscillations we subtracted the 1/f spectrum by multiplying the power spectrum with a quadratic model of the frequency spectrum (https://www.fieldtriptoolbox.org/example/ecog_ny/) which emphasizes the presence of narrowband peaks. Theta and Alpha frequency bands were defined based on the peaks of the 1/f background noise corrected power spectrum. The power spectrum was calculated by collapsing expected and unexpected conditions and hippocampal contacts using a multitaper frequency transformation with a Hanning taper from 1 to 30 Hz.

We calculated coherence for each patient, condition, and each pair of contacts. Given that coherence measures suffer from bias when comparing conditions with different number of trials (Bastos & Schoffelen, 2015), we randomly selected the number of trials for the expected condition to match the unexpected trial number, averaging across 500 randomizations for each subject. For each subject this yielded coherence and angle measures between all pairs of contacts, which were then averaged for all contacts in each portion (**Supplementary Table 2*Supplementary Table 2.*** ) at the theta and alpha frequency peak of the 1/f background noise corrected power spectrum. A repeated measures ANOVA with factors expectancy (expected, unexpected) *vs*. portion pairs (HH-HB, HH-HT, HB-HT) was performed for coherence across the 8 patients with contacts implanted in all hippocampal portions. Post-hoc comparisons between portion pairs were subsequently calculated using paired *t*-tests. For the coherence angle we used circular statistics (Fisher, 1995) provided by the MATLAB toolbox (CircStat2012a: https://es.mathworks.com/matlabcentral/fileexchange/10676-circular-statistics-toolbox--directional-statistics-) (Berens, 2009). A Harrison-Kanji test was used to calculate a circular two-factor ANOVA with factors expectancy (expected, unexpected) and inter-portion (HH-HB, HH-HT, HB-HT). For the post-hoc comparisons we used the Watson Williams test to test whether a set of mean angle differences between portions are equal.

#### Traveling waves

Theta oscillations have been shown to travel along the long axis of the hippocampus in humans (Zhang & Jacobs, 2015) and rodents (Patel et al., 2012). From our pool of 11 patients, we included 10 patients with consecutive electrodes along the long axis of the hippocampus and calculated traveling waves for the peak of the theta and alpha narrow band oscillations identified in the corrected power spectrum, using custom scripts running in Matlab. Trials were defined from -0.1 to 2 s peri-event onset. Since there were no differences in between-portion Coherency angle with novelty, traveling waves were calculated by collapsing across expected and unexpected conditions. Differences in the “traveling index” between expected and unexpected trials, and the modulation of direction of traveling (anterior to posterior vs posterior to anterior) by novelty, will be discussed elsewhere.

Next, we band pass filtered the data in the band of interest for each subject using a plateau shaped band-pass filter with a 15% transition zone. We chose the most anterior hippocampal contact as a reference channel and extracted 1s segments of the filtered data locked to the peaks of the reference channel (500 ms before and after the peaks of the reference channel) and averaged the extracted data per trial and per contact. The resulting band pass filtered signal locked to the reference channel was visually inspected for a traveling wave pattern in which amplitude peaks would advance (e.g., increase their peak time) from one contact to the next (**Supplementary Figure 4**). Next, we used a different approach to calculate traveling waves. In this approach, we first filtered the signal with the same plateau filter introduced before, and then we calculated the instantaneous phase via the angle of the Hilbert transform for each time point and each trial and contact. Using circular statistics (Fisher, 1995) within the MATLAB toolbox CircStat2012a (Berens, 2009) we calculated the circular distance between each contact and the reference - the most anterior contact - for every trial and averaged these circular distances by taking a circular mean across time points (1s from stimulus onset) for each trial. We then plotted an angle histogram across trials of the phase distance between every contact and the reference. A non-uniform distribution of the phase shift between each channel and the reference channel was tested using a Rayleigh test to assess phase locking across trials. The mean resultant vector length (phase locking value) and the angle were then calculated for the distance between the phase of each contact and the reference (most anterior) channel. In order to test whether every angle was significantly different from zero, we used a one sample test for the mean angle (the “mtest” function within CircStat2012a), similar to a one sample t-test on a linear scale (**Supplementary Figure 5**). We used this same procedure for comparing contact *vs*. reference channel, to calculate the phase distance between every pair of adjacent contacts (**Supplementary Figure 6**). The existence of a traveling wave along the long axis of the hippocampus would imply a constant phase distance between every pair of adjacent contacts. In order to define a traveling index, we pooled the phase distances between every pair of adjacent contacts and defined as traveling only those pooled angles significantly different from zero and non-uniform across trials, tested by means of a mtest and a Rayleigh test respectively.

Speed for the traveling waves was computed for each patient and each traveling wave frequency as:

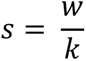

Where *w* is the angular frequency given by *w* = 2 *Πf*, *f* is the center frequency of the traveling wave and *k* is the wave number given by:

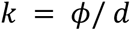

where *ϕ* is the absolute value of the phase distance between the most posterior electrode and the most anterior used in the traveling wave analysis and *d* the spatial distance between electrodes. Note that for **Figure 7** C and E, large phase jumps between electrodes in the hippocampal head or at the head-to-body transition, defined as the highest phase difference relative to the most anterior contact between every pair of contacts, were subtracted to compute *ϕ* for each patient and traveling wave frequency.

## Results

### Novelty-evoked responses. Intracranial event-related potentials to novelty show polarity inversion from HH to HB

Unexpected stimuli evoked a biphasic waveform following stimulus presentation in all three hippocampal portions (HH, HB and HT), with a first deflection peaking at ∼250ms followed by a second sustained deflection (**Figure 2A**). There was a polarity inversion of the iERP between the HH and the HB. That is, the first deflection was of negative polarity in the head but of positive polarity in the body and tail (**Figure 2A**). This polarity inversion was consistent across subjects; it was present in 11 of 11 patients and occurred approximately at the same latency for every patient (**Supplementary Figure 2**). For the 8 patients with electrode contacts in all hippocampal long-axis portions, a one way ANOVA on the differences between unexpected and expected conditions with within-subject factor portion (HH, HB, HT) revealed a significant interaction in an early time window (253-301 ms; p_clus_= 0.037; summed F values = 1.091 x 10^3^) (**Figure 2A** depicted in light grey, **Figure 2B**). Post-hoc *t*-tests on this early time window revealed that the interaction reflected HH and HB differences between expected and unexpected conditions (*t*_7_ = -4.837; p = 0.002). Furthermore, we found a significant difference between unexpected *vs*. expected condition (cluster-based corrected for multiple time points) at later latencies in the HB (555-572 ms; summed *t* values = 422.36; p = 0.02) and the HT (457-744 ms) (p = 0.0428; summed *t* values = -904.85) (**Figure 2A** depicted in dark grey).

**Figure 2.**
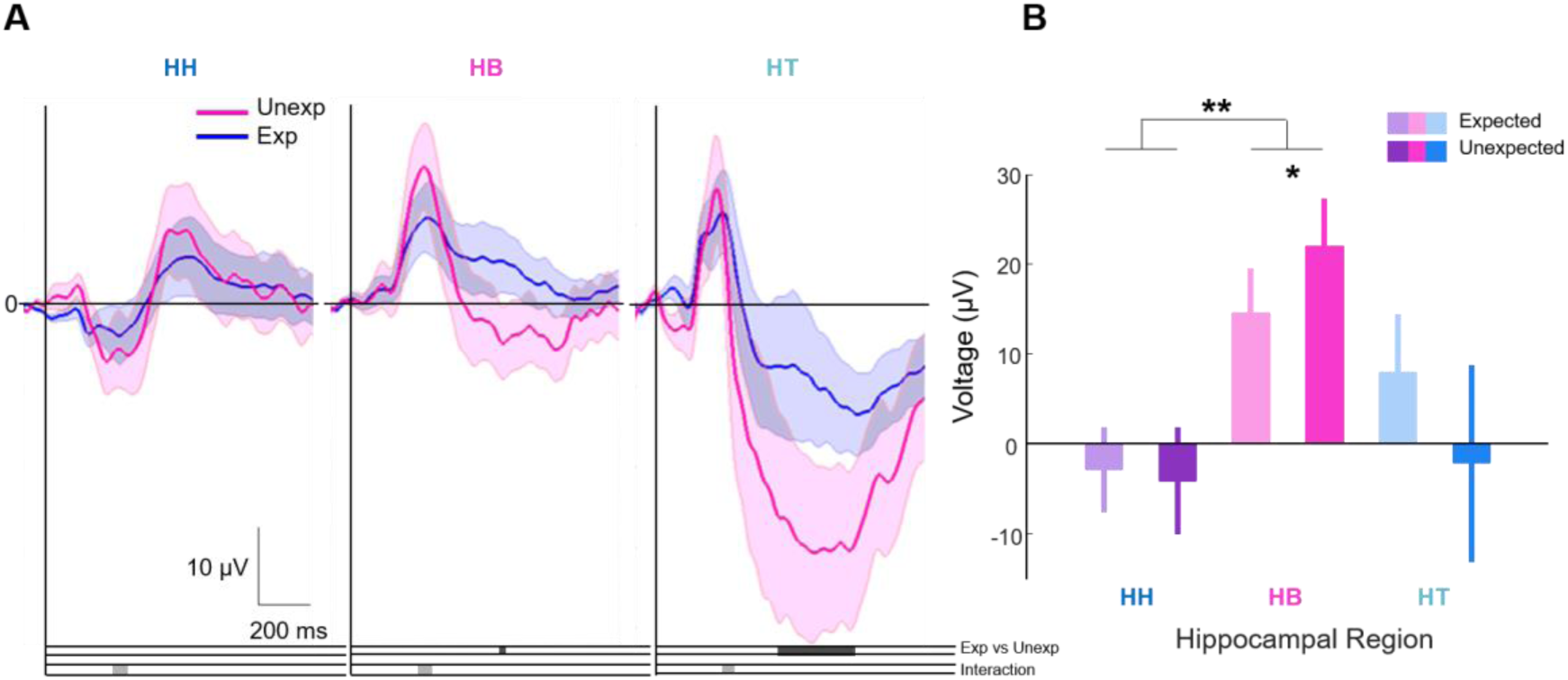
Hippocampal long-axis polarity inversion for intracranial event related potential (iERP) responses to unexpected stimuli. A. iERPs for expected (blue) and unexpected (red) trials, averaged across patients for each hippocampal long-axis portion separately. The interaction between conditions (unexpected and expected) and hippocampal portion (HH, HB, HT) in an early time window (253-301 ms) is indicated in light grey. Testing responses to expected *vs*. unexpected in individual portions revealed significant effects in the body (555-572 ms) and the tail (457-744 ms) depicted in dark grey. Negative potentials are plotted downwards; iERPs were referenced to an average reference across all hippocampal contacts. **B**. Averaged voltage in the time window in which the significant interaction between portion and novelty was found. Post-hoc *t*-test between conditions and portions reveal greater differences with novelty between the HH and HB. Error bars depict SEM. **(p<0.01), * (p<0.05).

### Novelty-induced responses. Power differences with novelty processing are higher in the hippocampal tail in the lower theta band

In a previous study, Axmacher et al. (2010) showed that theta (3–8 Hz) power increased at an early latency (200–400 ms) and later (500–1400ms) decreased during processing of unexpected as compared to expected items in the hippocampus. Higher gamma power (70–90 Hz) was selectively increased in the hippocampus during processing of unexpected items between 500–700 and between 1000–1100 ms (Axmacher et al. (2010)). Slow theta power increase has also been shown in the human hippocampus during associative prediction violations in a mismatch detection task (Chen et al., 2013). Here, using the same protocol as in (Axmacher et al., 2010) but specifically looking at the long-axis spatial localization of the hippocampus, we identified the same pattern of theta power increase and later decrease of the theta power in the comparison between unexpected and expected condition and an increase in gamma power in all 3 portions of the hippocampal long axis (**Figure 3, Supplementary Figure 3**). Specifically, cluster-based correction for multiple comparisons (time and frequency) between unexpected *vs.* expected items revealed a significant difference for the theta band in the HH (3-5Hz, 160-420ms, p_clus_ = 0.025, p_FDR_ = 0.025), in the HB (3-5Hz, 180-380ms, p_clus_ = 0.012, p_FDR_ = 0.014); and in the HT (1-11Hz, 0-450ms, p_clus_ = 0.001, p_FDR_ = 0.002). For the higher frequency range the cluster permutation test for the correction for multiple comparisons revealed a difference between expected and unexpected condition in the HH (60-90Hz, 120-750ms, p_clus_ = 0.003, p_FDR_ = 0.005), in the HB (67-112Hz, 60-800ms, p_clus_ = 0.001, p_FDR_ = 0.002), and in the HT (80-150Hz, 90-1000ms, p_clus_ = 0.001, p_FDR_ = 0.002) (**Figure 3 A, Supplementary Table 3.,** False Discovery Rate, FDR, correction applied to the 6 p values for theta and gamma clusters). Note that, although the significant gamma activity cluster frequency appears to increase from anterior to posterior, taking frequency centre of mass we did not find a significant change with long-axis position, either as a linear or a step-wise change.

**Figure 3.**
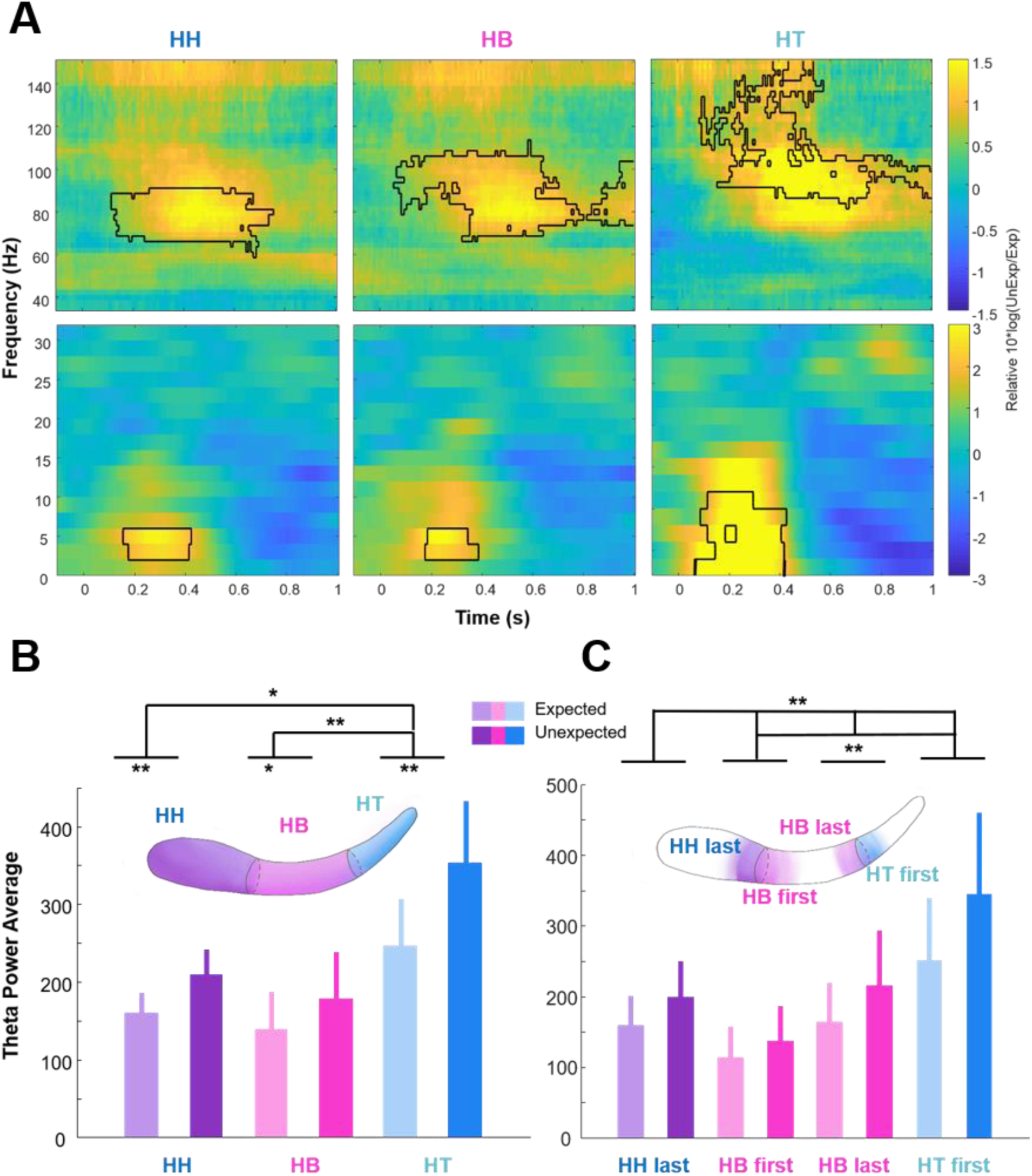
Gradual increase in theta power along the hippocampal long axis in response to novel stimuli A. Similar oscillatory responses to novelty across long-axis portions of the human hippocampus. Time-frequency plots for the logarithmic relative change in power between expected and unexpected conditions. *T*-tests between expected and unexpected conditions were performed separately within each portion (HH, HB and HT in columns) and for lower (0-33Hz) frequencies (lower row) and higher (34-150Hz) frequencies (upper row). Significant clusters that survived correction for multiple comparisons are outlined in black. **B-C. Distinct oscillatory responses to novelty across long-axis portions of the human hippocampus. B.** Post-hoc *t*-tests on the averaged power in the significant time frequency window from the cluster-based permutation test for the one way ANOVA on the differences between unexpected and expected condition *vs*. portion (HH, HB and HT). Post-hoc *t*-tests revealed greater theta-power differences between unexpected and expected stimuli in the tail compared to differences in HB and HH. **C.** Same as B. averaging across first and last contact of each region for each patient shows a gradual increase in theta power from HB to HT. Error bars depicts the SEM. **(p<0.01), * (p<0.05)

We next selected an *a priori* time window (250-400ms) of interest in our time frequency data based on previous findings attributing a novelty effect in the theta and slow-theta frequency ranges (Axmacher et al., 2010; Chen et al., 2013). The cluster-based permutation tests over low frequencies (1-20 Hz) in this time window revealed a significant interaction between expected and unexpected conditions across the different portions of the hippocampus (HH, HB and HT) from 1 to 5 Hz (p_clus_ = 0.003; summed F values = 63.23). A one way ANOVA on the differences between expected and unexpected condition with within-subject factor portions (HH, HB, HT) on the average power in this time window (1-5 Hz; 250-400 ms), revealed a significant main effect of portion (F(1.7,11.93) = 6.44; p = 0.01), a significant main effect of novelty (F(1, 7) = 23.07; p = 0.02), and a significant interaction between region and novelty (F(1.14, 7.98) = 8.13; p = 0.02). Post-hoc *t*-tests on the average power in the significant time frequency window revealed a higher difference between unexpected and expected conditions in the HT as compared to HB and HT. The exact statistics are as follows: HT expected vs. unexpected (1-5 Hz; 250-400 ms, t(7) = -4.57, p = 0.003); HB expected vs. unexpected (1-5 Hz; 250-400 ms, t(7) = -2.72, p = 0.03); HH expected vs. unexpected (1-5 Hz; 250-400 ms, t(7) = -4.3, p = 0.004). Post-hoc *t*-tests on the differences between expected and unexpected condition inter-region also suggested that differences were most pronounced in HT: HHdiff vs HBdiff (1-5 Hz; 250-400 ms, t(7) = 1.24, p = 0.25); HBdiff vs HTdiff (1-5 Hz; 250-400 ms, t(7) = -3.78, p = 0.07); HHdiff vs HTdiff (1-5 Hz; 250-400 ms, t(7) = -2.37, p = 0.05); (**Figure 3B**).

This time-frequency analysis revealed differences in novelty processing along the long axis of the hippocampus in a low frequency range (1-5 Hz) showing greater differences between the tail and the rest of the hippocampus. Notably, theta power in the hippocampal body was at an intermediate level between head and tail, *i.e.*, the change in theta power appeared as a gradual increase from body to tail (**Figure 3C**).

In a similar analysis, taking an *a priori* time-frequency window during which gamma responses to unexpected stimuli have been shown to occur (40-150 Hz averaging between 500–700 ms (Axmacher et al. (2010)), we found no interaction with hippocampal portion (non-significant cluster at frequency 100Hz, p_clus_ = 0.407; summed F value = 13.83). This demonstrates that novelty-related power differences were specific to the theta-frequency range.

### Higher alpha and theta coherence between posterior and anterior hippocampus with higher offsets between HH and HB regardless of expectancy

We next tested whether oscillations in different long-axis portions became more or less synchronized as a function of expectancy. Given that our findings revealed differences along the long axis for the lower frequencies (time-domain evoked responses and low-frequency induced power responses), together with the established roles for theta/alpha rhythms in synchronization properties (Buzsáki, 2002; O’keefe & Nadel, 1978; Zhang, Watrous, Patel, & Jacobs, 2017), we focused on low-frequency bands for subsequent coherence and phase analyses. Hence, we tested whether inter-portion coherences in the theta/alpha bands varied between expected and unexpected items. For this we computed the absolute value (Coherence) and the angle (Phase Angle) of the complex Coherency measure between every pair of contacts. For each frequency band, a repeated measures ANOVA with factors inter-portion (HH-HB, HH-HT, HB-HT) and Expectancy (expected, unexpected) on the absolute value of the coherency (Coherence) revealed a main effect of Inter-portion in both frequency bands (theta: F(2,14) = 3.518; p < 0.05; alpha: F(2,14) = 7.123; p < 0.007) (**Figure 4A**). We did not observe an interaction between inter-portion and expectancy in either frequency band (theta: F(2,14) = 0.451; p = 0.646; alpha: F(2,14) = 1.568; p = 0.243). The post-hoc t-tests – averaging coherence values for expected and unexpected conditions – revealed a difference between HH-HT and HB-HT coherence, being higher for HH-HT, reflecting less variability between subjects in the coherency angle (**Figure 4B**) (theta: t_7_ = 2.311, p = 0.05; alpha: t_7_ = 4.4; p = 0.003) (**Figure 4A**). Given that rat CA1 theta coherence diminishes from dorsal to ventral hippocampus (corresponding to human posterior to anterior hippocampus) (Patel et al., 2012), this observation of more coherence between posterior and anterior hippocampal portions than with the middle portion is striking and unexpected.

**Figure 4.**
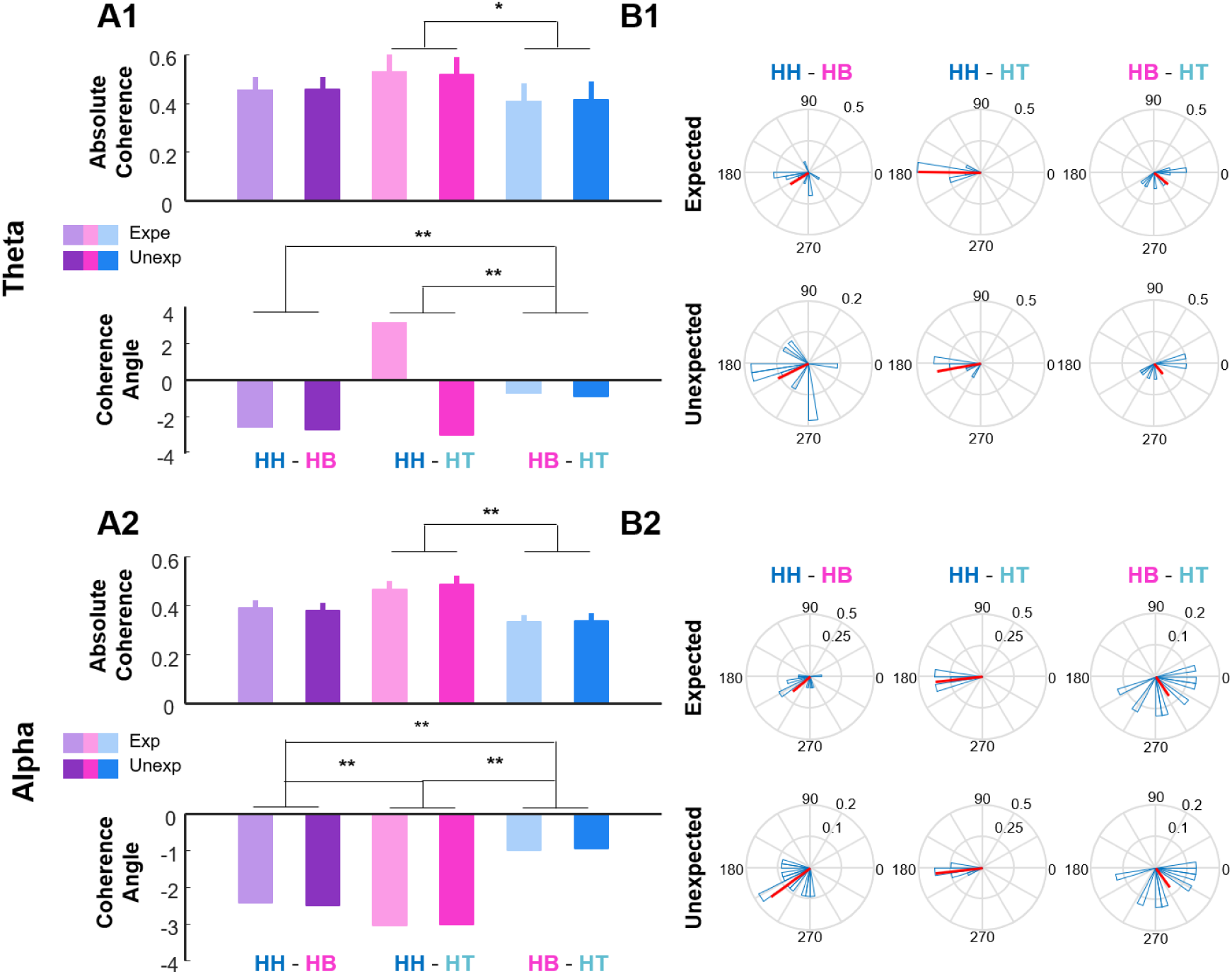
Theta and alpha coherence transition along the long-axis, independently of stimulus expectancy. **A.** Between-portion absolute coherence averaged over patients for theta (A1) and alpha (A2) oscillations. Error bars show SEM. A repeated measures ANOVA with factors Inter-portion (HH-HB, HH-HT, HB-HT) and Expectancy (expected, unexpected) on the absolute value of the coherence revealed a main effect of Inter-portion. Coherence Angle: A circular two-factor ANOVA with factors Inter-portion (HH-HB, HH-HT, HB-HT) and Expectancy (expected, unexpected) on the angle of the coherence revealed a main effect of Inter-portion. Results of post-hoc t-tests are shown. **B.** Polar histograms across patients for the coherency (angle and module), circular mean (depicted in red) across portions and patients (depicted in blue). **(p<0.01), * (p<0.05)

We next performed a similar analysis on the coherency angle, using a circular two-factor ANOVA with factors Inter-portion (HH-HB, HH-HT, HB-HT) and Expectancy (expected, unexpected). This analysis again revealed a main effect of Inter-portion (theta: df = 4, χ = 38.74; p < 7.87 10^-8^; alpha: df = 4, χ = 49.35; p < 4.93 10^-10^). The post-hoc *Watson William*-tests showed significant differences between the circularly averaged angle differences for expected and unexpected conditions between HH-HB *vs*. HB-HT (theta: F_1,14_ = 9.64; p = 0.008; alpha: F_1,14_ = 10.05; p = 0.007), HH-HT *vs*. HB-HT (theta: F_1,14_ = 21.98; p = 3.49 x 10^-4^; alpha: F_1,14_ = 27.10; p = 1.33 x 10^-4^) and HH-HB *vs*. HH-HT for alpha (F_1,14_ = 7.92; p = 0.01) only (theta: p = 0.22). This result indicated a cumulative phase lag of 180° between anterior and posterior portions of the hippocampus as has been previously suggested in rodents (Patel et al., 2012). However, whereas in the rat CA1 the phase advance is monotonic with dorsal to ventral distance (Patel et al., 2012), we show a monotonic phase advance along the long axis with discrete phase offset between the HH and HB.

### Traveling waves in the theta and alpha band as a common processing mechanism

We have shown that coherence angle between HH-HT is *∼pi* radians which potentially reflects the summation of coherence phases from HH-HB and HB-HT. Given the possible cumulative phase advance observed in the Coherency angle along the long axis, we next aimed to test for the existence of previously suggested traveling waves (Patel et al., 2012; Zhang & Jacobs, 2015) along the hippocampal long axis in the theta and alpha bands. In view of the lack of a modulation of phase advance by novelty, we tested for long-axis traveling waves regardless of novelty. This was done for theta as well as for the alpha band, defining our frequency bands based on the peaks for narrow-band oscillations found in the 1/f corrected power spectrum (**Table 1**). The theta traveling waves thus far demonstrated in humans used a broad frequency band (2-10 Hz) (Zhang & Jacobs, 2015). Here we tested whether distinct alpha peaks in the power spectrum corresponded to distinct alpha traveling waves, akin to those observed in the human neocortex (Bahramisharif et al., 2013; Zhang et al., 2017).

**Table 1.** Frequency ranges for the theta and alpha bands defined from the frequency peaks in the 1/f corrected power spectrum. In bold: not traveling (patient 2 in the alpha band, patient 6 in theta band) for the patients included in the traveling wave analysis.

| Patient | $\Theta$ | | | $\alpha$ | | |
| --- | --- | --- | --- | --- | --- | --- |
|  | Range | Middle freq | peak | Range | Middle freq | peak |
| **2** | 2-6 | 4 | 3 | **12-14** | **13** | **13.3** |
| **3** | 3-5 | 4 | 4.3 | 14-16 | 15 | 14.8 |
| **4** | 2-4 | 3 | 2.4 | 13-15 | 14 | 14.3 |
| **5** | 2-6 | 4 | 4.3 | 12-14 | 13 | 12.9 |
| **6** | **2-6** | **4** | **3** | 10-12 | 11 | 11.0 |
| **7** | 7.5-9.5 | 8.5 | 8 | 12-15 | 13 | 13.3 |
| **8** | 7-9 | 8 | 7.6 | 12-16 | 14 | 13.8 |
| **9** | 4-6 | 5 | 5.2 | 9-11 | 10 | 10 |
| **10** | 5-7 | 6 | 5.7 | 8.5-10.5 | 9.5 | 9 |
| **11** | 2-6 | 4 | 3 | 8-12 | 10 | 9.5 |

**Table 2.** Traveling Index for the theta band. Rayleigh test and mtest for the pooled angles between adjacent contacts. h = 0 if null hypothesis cannot be rejected, 1 otherwise; upperlc and lowerlc: upper and lower confidence level respectively.

| patient | $\theta$ | | | | | |
| --- | --- | --- | --- | --- | --- | --- |
|  | Rayleigh test |  |  | mtest |  |  |
|  | p | z | h | mean | upperlc | lowerlc |
| 2 | $1.82 \cdot 10^{-297}$ | 621.20 | 1 | -0,06 | -0,01 | -0,11 |
| 3 | $3.01 \cdot 10^{-297}$ | 626.14 | 1 | -0,16 | -0,11 | -0,21 |
| 4 | 0 | 982.98 | 1 | 0,14 | 0,18 | 0,10 |
| 5 | $1.88 \cdot 10^{-53}$ | 115.81 | 1 | 0,34 | 0,46 | 0,21 |
| 6 | $1.078 \cdot 10^{-14}$ | 31.83 | 0 | 0,10 | 0,35 | -0,14 |
| 7 | 0 | 889.45 | 1 | -0,19 | -0,15 | -0,23 |
| 8 | 0 | $1.04 \cdot 10^3$ | 1 | -0,08 | -0,04 | -0,12 |
| 9 | $1.30 \cdot 10^{-293}$ | 616.99 | 1 | -0,27 | -0,21 | -0,32 |
| 10 | $4.95 \cdot 10^{-136}$ | 288.00 | 1 | -0,15 | -0,08 | -0,23 |
| 11 | 0 | $1.40 \cdot 10^3$ | 1 | -0,17 | -0,14 | -0,21 |

We observed the theta and alpha peaks of the LFP signals (locked to the most anterior hippocampal contact as a reference) advancing in time from posterior to anterior contacts along the long-axis of the hippocampus as has been previously reported in a broader frequency band (2-10 Hz), centered on theta but encroaching into delta and alpha bands (Zhang & Jacobs, 2015). Representative patients illustrate the theta and alpha peaks in their power spectrum (**Figure 5 1 A, 2 A**), electrode contact positions (**Figure 5 1 B, 2B**) and traveling wave pattern (**Figure 5 1C, 2C**). The cumulative phase increase observed for both theta and alpha rhythms is in agreement with the existence of traveling waves along the long axis of the hippocampus reported in rodents (Lubenov & Siapas, 2009; Patel et al., 2012) and in humans (Zhang & Jacobs, 2015)

**Figure 5.**
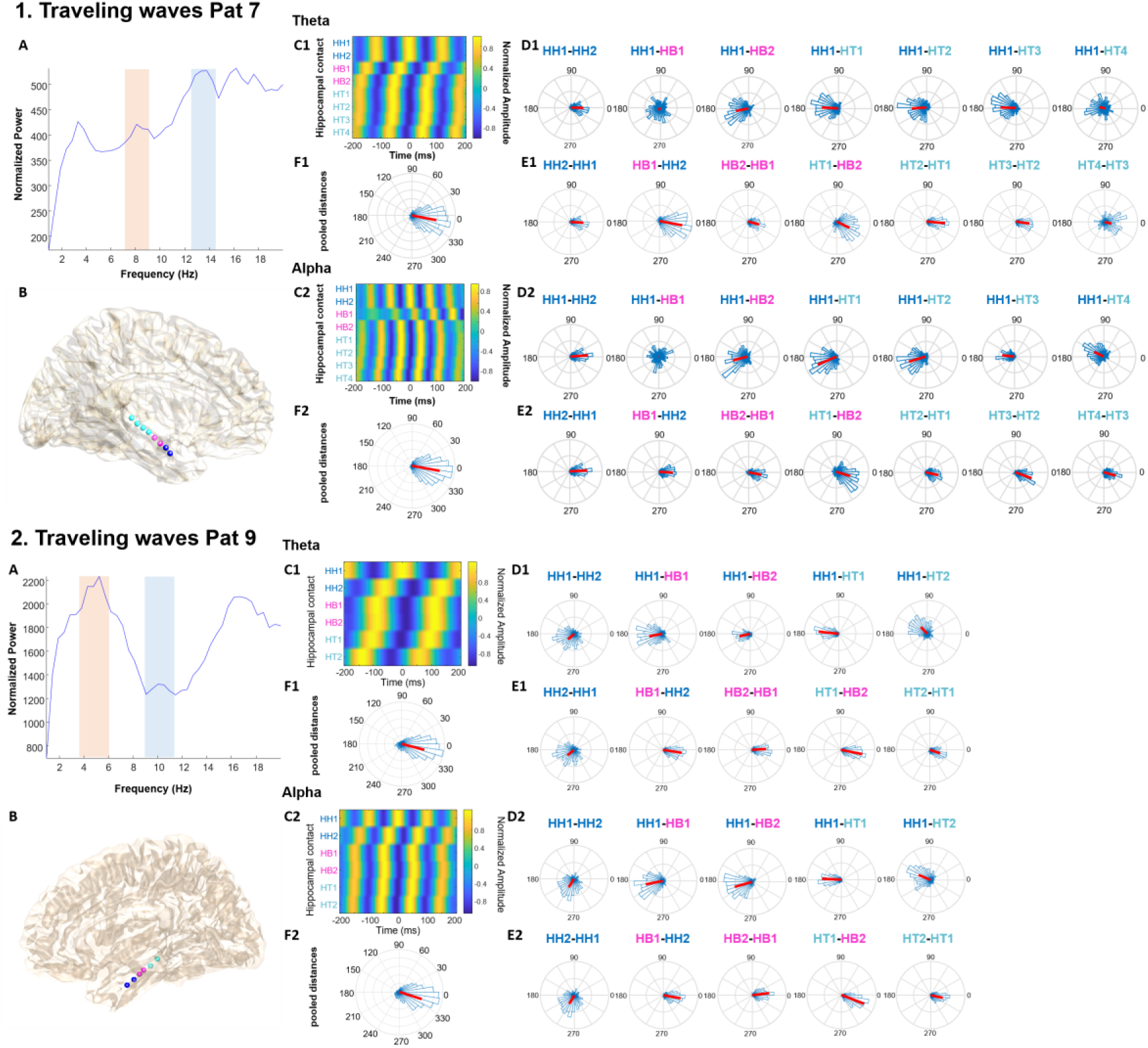
1. Theta and alpha traveling waves for Patient 7. A. power spectrum collapsing across hippocampal electrodes with 1/f background signal correction. Pink and blue shadings depict the frequency bands used for the definition of the theta and alpha bands, respectively. No traveling waves were found for this patient in the slower theta band. **B**. Brain image depicting the locations of depth electrodes implanted in Patient 7. Hippocampal contacts in head, body and tail are depicted schematically in dark blue, pink and turquoise, respectively on an anatomical template in MNI space. (see **Supplementary Figure1** for Pre- and post-implantation MRI for each patient) **C**. LFP peaks advancing from posterior to anterior portions of the hippocampus. Filtered signals locked to the peaks (yellow) of the most anterior hippocampal electrode (HH1) as a reference channel and averaged across trials. Effects are shown for theta (C1) and alpha (C2) oscillations. **D**. Polar histogram across trials for phase differences between the most anterior electrode as a reference channel and each electrode along the longitudinal axis of the hippocampus. **E**. Polar histogram across trials for the phase difference between every pair of adjacent channels along the longitudinal axis of the hippocampus. **F**. Polar histogram across trials for the pooled phase differences between all pairs of adjacent channels along the longitudinal axis of the hippocampus as a traveling index. **2. Traveling waves for Patient 9 (as for Figure 5.1).**

In our next analysis, a pattern of activity advancing in time was assessed by analyzing the instantaneous phase in every contact and comparing phase distances between each contact and the most anterior contact as a reference (**Figure 5 1 D**, **2 D**). Rayleigh tests revealed that the phase distance between each channel and the reference channel was different from a uniform distribution for the majority of the contacts. Furthermore, a one sample m-test (one-sample test for the mean angle) revealed phase angles between the majority of the contacts and the reference were different from zero for theta and alpha bands (**Supplementary Table 4.** ). As another measure, we also assessed non-uniform and different-from-zero phase differences between pairs of adjacent contacts (***Figure 5 1 E, 2 E, Supplementary Table 5. , Supplementary Table 7.***). We found these to be generally lower in posterior portions of the hippocampus compared to phase differences observed in anterior portions, in line with changes in Coherency from posterior to anterior (as described above).

Furthermore, in order to provide a “traveling index” we assessed non-uniform and different-from-zero pooled phase distances between every pair of adjacent contacts for the theta and alpha bands (**Figure 5 1 F, 2 F, Table 1, Table 3, Supplementary Table 5. , Supplementary Table 7.** ). As expected from the results of our coherency analysis, higher phase differences were observed in the anterior hippocampus. This abrupt dichotomy of phase difference for the theta and alpha bands, being larger in the anterior portion of the hippocampus and more constant between adjacent contacts in the posterior portion (closer to body and tail), mirrors the Coherency angle differences observed between portions (**Figure 4**) and the iERP reversal (**Figure 2**). In line with the coherency angle results (**Figure 5**), the traveling wave analysis also revealed that theta and alpha oscillations were in antiphase (∼180°) between anterior and posterior portions in the majority of patients, as has been previously reported in rodents (Patel et al., 2012).

**Table 3.** Traveling Index for the alpha band. Rayleigh test and mtest for the pooled angles between adjacent contacts. h = 0 if null hypothesis cannot be rejected, 1 otherwise; upperlc and lowerlc: upper and lower confidence level respectively.

| patient | $\alpha$ | | | | | |
| --- | --- | --- | --- | --- | --- | --- |
|  | Rayleigh test |  |  | mtest |  |  |
|  | p | z | h | mean | upperlc | lowerlc |
| 2 | $4.07 \cdot 10^{-281}$ | 590.21 | 0 | -0,03 | 0,02 | -0,08 |
| 3 | $4.30 \cdot 10^{-314}$ | 658.30 | 1 | -0,26 | -0,21 | -0,31 |
| 4 | 0 | $1.18 \cdot 10^3$ | 1 | 0,063 | 0,10 | 0,03 |
| 5 | $5.18 \cdot 10^{-35}$ | 76.60 | 1 | -0,42 | -0,26 | -0,57 |
| 6 | $1.13 \cdot 10^{-19}$ | 43.01 | 1 | 0,21 | 0,42 | 0,0001 |
| 7 | 0 | $1.08 \cdot 10^3$ | 1 | -0,19 | -0,15 | -0,23 |
| 8 | 0 | $1.16 \cdot 10^3$ | 1 | -0,13 | -0,09 | -0,17 |
| 9 | 0 | 681.13 | 1 | -0,32 | -0,27 | -0,37 |
| 10 | $2.66 \cdot 10^{-165}$ | 344.07 | 1 | -0,11 | -0,04 | -0,18 |
| 11 | 0 | $1.45 \cdot 10^3$ | 1 | -0,23 | -0,20 | -0,26 |

### Traveling waves in the theta and alpha bands: backwards or forwards with phase jumps in the anterior hippocampal portion

Closer inspection of the phase progression through the hippocampal long axis (**Figure 6**) revealed several important properties. Firstly, across different patients, waves travelled in either the anterior-to-posterior (positive phase distance) and posterior-to-anterior (negative phase distance) direction, with the latter direction being more frequently observed (Zhang & Jacobs, 2015; Zhang, Watrous, Patel, & Jacobs, 2018). Secondly, the direction of travel for theta *vs*. alpha waves were found to be different in the same patient in 3 cases (compare patients 5, 7 and 11 in **Figure 6A** *vs*. **Figure 6B**). Thirdly, across all patients, there were clear phase jumps (of up to ∼3 radians) in the anterior portion, commonly at the HH to HB transition. This was evident for both theta and alpha oscillations and occurred irrespective of the direction of travel. Control analyses suggested that our effects are unlikely to be due to transitions between hippocampal subfields (see Supplementary Results),

**Figure 6.**
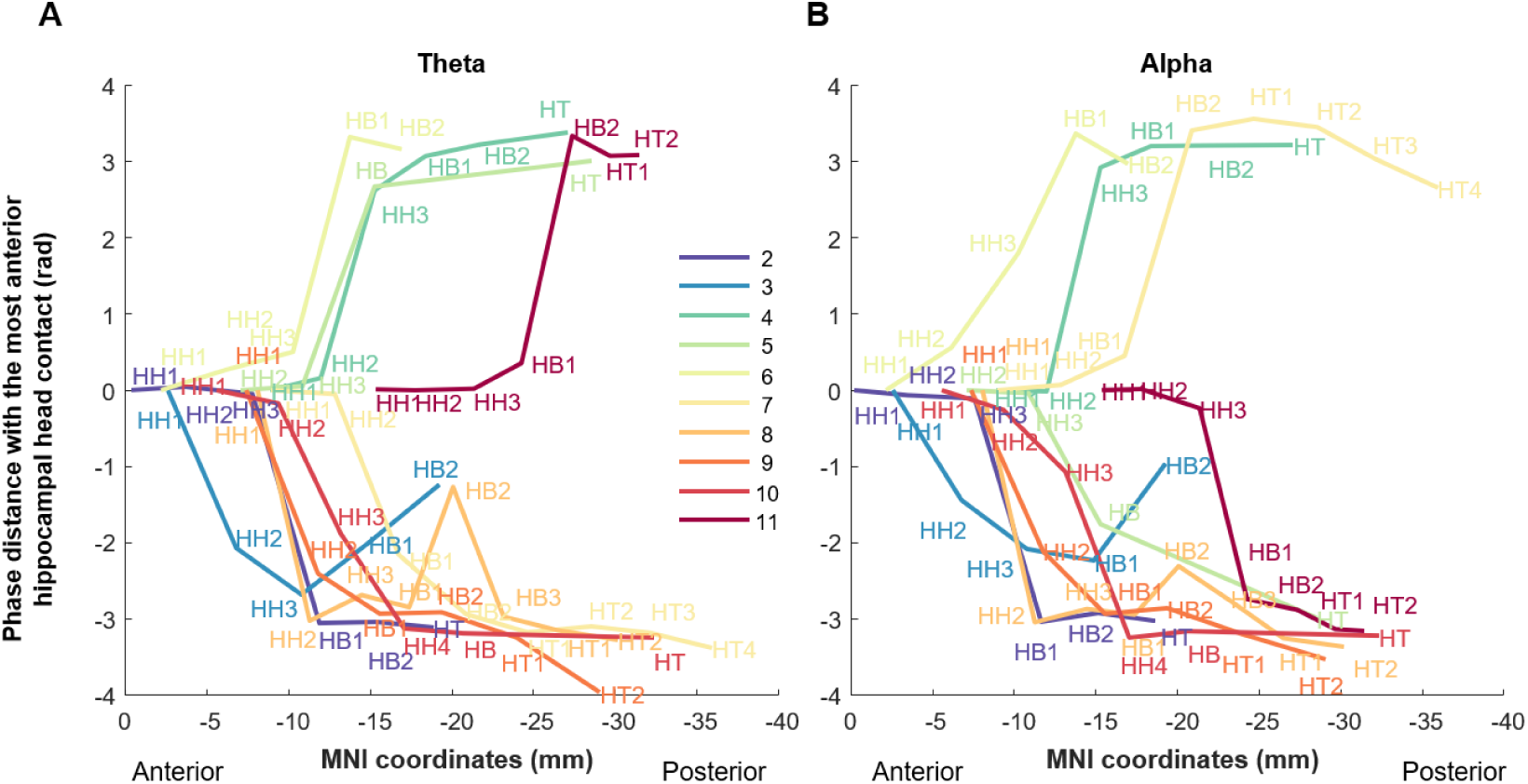
Hippocampal traveling waves travel in both posterior-to-anterior and anterior-to-posterior directions, showing marked phase jumps at the transition between head and body. Phase distance for theta (**A**) and alpha (**B**) traveling waves are plotted across the hippocampal long axis, relative to the most anterior hippocampal contact for every patient. Each hippocampal contact is labeled, with color distinguishing electrodes from separate patients (legend: patient number and color). The horizontal axis indicates the contact’s distance to the anterior tip of hippocampus in MNI space (the *y* coordinate). The vertical axis indicates the contact’s estimated unwrapped phase distance relative to the most anterior contact of the hippocampus.

We next proceeded to measure the speed of traveling wave propagation for traveling waves for each patient. The distribution of individual traveling wave frequencies in the theta (∼3-8 Hz) and alpha (∼9-15 Hz) bands is shown in **Figure 7A** (corresponding to middle frequency in **Table 1**). Traveling speed varied by an order of magnitude depending on whether the large phase jumps were included **Figure 7**) or excluded (**Figure 6C**) from the calculation. When excluding large phase jumps, phase differences between the most anterior and most posterior contacts used to compute *k* decreased so that speed reached similar values as found in a previous study (Zhang & Jacobs, 2015) where no such large phase jumps were reported. In both cases, speed increased monotonically with increasing frequency (**Figure 7D-E**), irrespective of the direction of travel (Pearson’s r = 0.81, P < 0.00001 with phase jumps included; Pearson’s r = 0.47, P < 0.03 excluding phase jumps).

**Figure 7.**
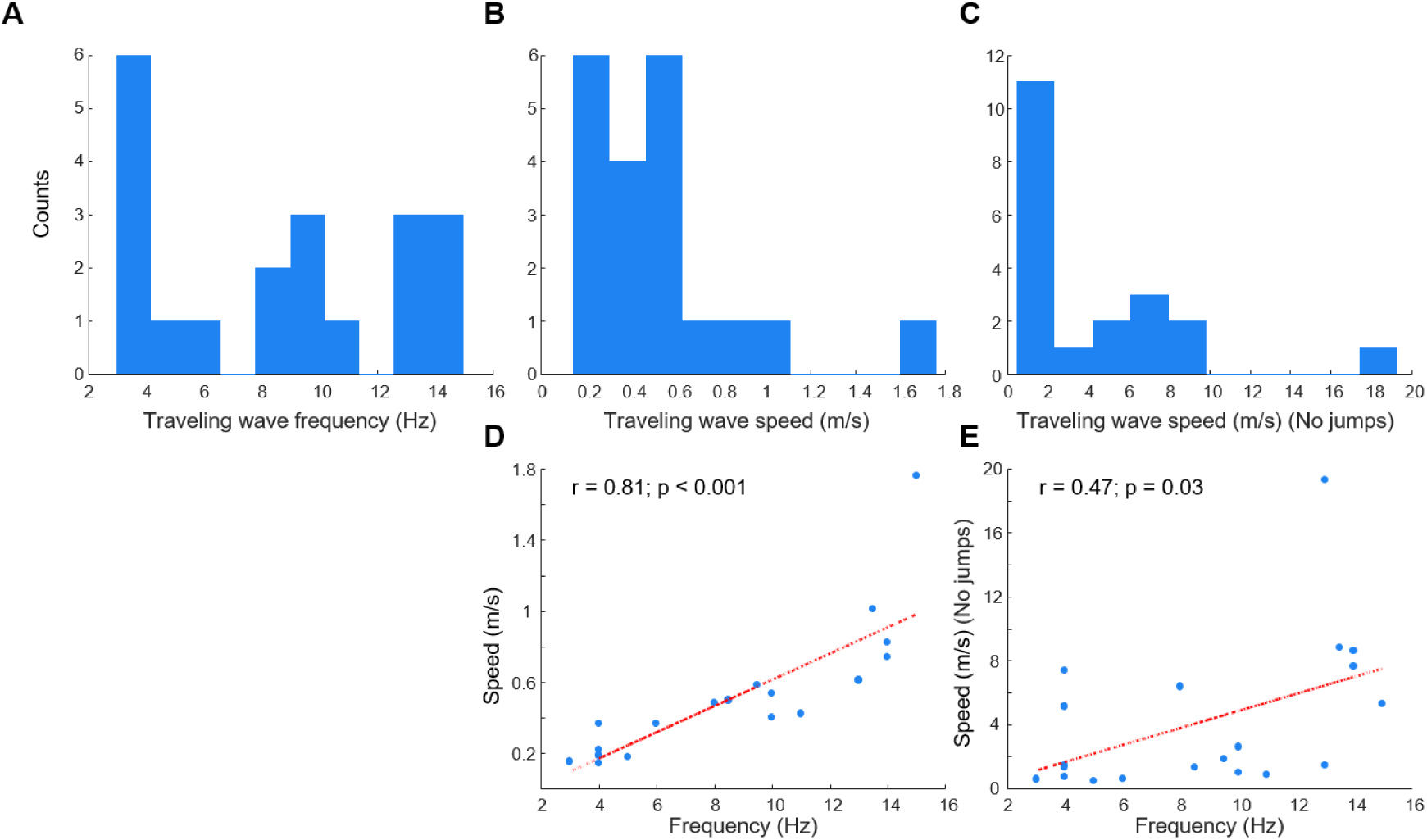
Population analysis of hippocampal traveling waves for alpha and theta frequencies. Histograms of the frequency distribution of the theta and alpha traveling waves (**A**), the speed for each traveling wave (**B**), and the speed for each traveling wave (**C**) removing large phase jumps observed in the hippocampal head or head-to-body transition. **D**. Speed as a function of traveling wave frequency. Red line indicates a best fit regression line, inset indicates correlation coefficient and P value. **E**. Same as D but removing large phase jumps observed in the hippocampal head or head-to-body transition.

## Discussion

A novel avenue for understanding hippocampal function is studying the different roles of portions along the hippocampal long axis (Brunec et al., 2018; Collin et al., 2017; Moser & Moser, 1998; Poppenk et al., 2013; Schlichting et al., 2015; Strange et al., 2014; B. Strange et al., 1999; Zeidman & Maguire, 2016). The current results provide several new insights into the hippocampal long axis novelty response. Time domain responses (iERPs) showed a novelty-evoked response along this axis but with a polarity inversion between the HH and HB in all patients, potentially indicative of a local generator at this junction. In the time-frequency domain, novelty-induced activity again suggested the entire hippocampal long axis to be involved in novelty processing, with all 3 portions showing a theta and gamma power increase. However, novelty effects on theta oscillatory power showed a different long-axis pattern as compared to iERPs: a gradual increase from HB to HT without any sharp transition. Independent of novelty, between-portion theta (and alpha) coherence dropped between HH and HB, with traveling waves displaying an abrupt change in phase at the same anatomical location.

### iERP polarity inversion between HH and HB

Locally generated large amplitude human hippocampal responses to unexpected stimuli are a consistent observation in intracranial recordings (Barbeau et al., 2017; Brázdil et al., 2001; Fell et al., 2005; Grunwald et al., 1999; Halgren, 1986; Halgren et al., 1995; Halgren et al., 1980; Ludowig, Bien, Elger, & Rosburg, 2010; McCarthy et al., 1989; Roman, Brázdil, Jurák, Rektor, & Kukleta, 2005; Smith et al., 1990; Stapleton & Halgren, 1987). Here, however, we demonstrate a reliable and marked polarity inversion within the hippocampus itself. Such a finding is crucial for undertanding the hippocampal role in novelty processing as it provides evidence that the actual generator of the MTL-P300 lies within the hippocampus (Mitzdorf, 1985). Our results are in line with pioneering studies on patients with longitudinally implanted electrodes demonstrating polarity inversions for oddball-evoked iERPs from anterior to porterior hippocampus in individual patients (Brázdil et al., 2001; McCarthy et al., 1989; Rosburg et al., 2007; Smith et al., 1990). However, the interpretation of these previous findings was limited due to imprecision because of the absence of MRI scans for electrode localization (Brázdil et al., 2001; McCarthy et al., 1989; Rosburg et al., 2007; Smith et al., 1990). In contrast, as one of several new insights provided here, we are the first to demonstrate such polarity inversion within the hippocampus using precisely localized electrode positions.

A more recent study on oddball-evoked hippocampal responses, measured in patients with longitudinally implanted electrodes, showed a linear increase of MTL-P300 mean amplitudes along the longitudinal axis of the HB, with largest amplitudes in posterior HB. In the subiculum, larger MTL-P300 mean amplitudes were found more anteriorly (Ludowig et al., 2010). In that study, however, depth electroencephalograms were referenced against linked mastoids, thus polarity inversions along the long-axis could not be measured. By contrast, in the current study, we demonstrated a polarity inversion of the oddball-evoked iERP within the hippocampus consistently at the same anatomical locus in all patients, thereby indicating a source for the novelty signal between hippocampal head and body.

### Novelty-induced theta power is highest in the hippocampal tail

Theta and gamma power increased in response to unexpected stimuli in all long-axis portions of the hippocampus. This accords with previous findings in rodents (Penley et al., 2013) where locomotion in a novel environment was related to increases in theta and gamma power at most CA1 and DG sites. A recent human intracranial study reported different theta frequencies along the long-axis, being slower in anterior (3 Hz) than posterior hippocampus (8 Hz), depending on movement speed in the virtual environment (Goyal et al., 2020). The novelty-induced theta power increases we observed overlapped in their frequency, with significant effects in HH/HB at 3-5 Hz and HT 1-11 Hz. We did not, however, find significantly greater power in posterior hippocampus (HT) *vs*. head and body for the non-overlapping higher theta range (5-11 Hz). The power increase with novelty processing was higher for HT compared to HH and HB in the slow-theta frequency range (1-5 Hz). These slow theta band power increases in posterior *vs*. anterior hippocampus have been shown in rodents (dorsal greater than ventral) during spatial navigation (Royer, Sirota, Patel, & Buzsáki, 2010) and to predict successful episodic memory encoding in humans (Lin et al., 2017). Thus, this increase in the slow theta band posteriorly for novelty processing may possibly relate to memory encoding of unexpected events.

### Abrupt drop in theta and alpha coherence across the hippocampal long axis

In rats, theta-wave coherence is relatively high between dorsal and intermediate hippocampal portions (equivalent to tail and body) but substantially lower between dorsal and ventral portions (equivalent to tail and head) (Patel et al., 2012). However, in response to a novel spatial context, theta coherence in CA1 has been shown to increase between dorsal and ventral poles more than between dorsal and intermediate portions (Penley et al., 2013). By contrast, our findings in humans suggest higher coherence between anterior (HH) and posterior portions (HT) compared to adjacent portions of the hippocampus (*e.g.*, HB-HT), but without a modulation by novelty. The higher coherence between HH and HT was accompanied by greater phase coherence offsets between HH and HB.

### Traveling theta and alpha waves

Traveling waves have been observed along the long axis of the hippocampus in the theta band in rodents traveling in the dorsal to ventral direction (Lubenov & Siapas, 2009; Patel et al., 2012). In humans, traveling waves have been reported transitioning from posterior to anterior hippocampus (dorsal to ventral in rodents) at a broader range of frequencies (2-10 Hz) compared with rodents (Zhang & Jacobs, 2015). This cross-species evidence highlights the potential functional importance of traveling hippocampal theta oscillations, possibly via the coordination of phase coding throughout the hippocampus in a consistent manner. That is, traveling waves engender that, at a given moment, there are gradually changing phases of the theta signal in different longitudinal portions of the hippocampus, which might represent a mechanism to code information within the hippocampus and in concert with cortical areas.

Concomitant with stronger coherence between opposites poles of the hippocampal formation along the long axis, a cumulative coherence phase increase from posterior to anterior of approximately -180° was found, in line with rodents studies (Patel et al., 2012), There was a more monotonic phase distance between pair of adjacent contacts within posterior (HT) and intermediate (HB) hippocampal portions, whereas a higher phase offset was found between anterior (HH) and intermediate portions. The latter was not reported in rodent recordings. This higher phase offset between anterior and intermediate portions is in line with the greater coherence angle offsets found in the same frequency bands between HH and HB. The fact that in previous studies in humans this phase offset was not observed could be due to the clustering of contacts used and the exclusion criteria for the contacts in the clusters (Zhang & Jacobs, 2015), perhaps leading to a higher exclusion of anterior electrodes which have been shown here to present the higher phase differences relative to the rest of the hippocampus.

Given previous cross-species evidence that theta waves travel in the dorsal-ventral (posterior-anterior) direction, our finding that traveling waves in some patients moved in the opposite (anterior-posterior) direction was unexpected, although we note recent evidence for bidirectional traveling waves recorded from a limited portion of the long-axis using thin-film microgrid arrays conformed to the human hippocampal surface (Kleen et al., 2021). This observation that theta and alpha traveling waves may, in the same patient, be traveling in opposite directions, suggests further flexibility in the routing of different information types with cortical areas. Regardless of direction of travel, speed increased with frequency indicating a common mechanism of propagation in either direction. The frequency-dependence of wave propagation speeds, as opposed to a constant speed for all frequencies, has been interpreted as weakly coupled intrahippocampal-entorhinal matched oscillators generating theta traveling waves (Kopell & Ermentrout, 1986; Patel et al., 2012; Zhang & Jacobs, 2015). The current results support these previous findings, by showing a monotonic increase in speed with increasing frequency and extend these previous findings to traveling waves in the alpha range.

### The hippocampal head to body transition

In addition to a change in polarity of novelty-evoked ERP at the HH-HB junction, we show a marked change in the phase of coherence and phase shift in the long-axis traveling wave at the same anatomical locus for alpha and theta oscillations. These findings are in accordance with a previous study, with measurements at adjacent contacts with the same electrode type as described here, showing an abrupt decrease in coherence at approximately the transition between the anterior one-third and posterior two-thirds of the hippocampus (Staresina, Fell, Do Lam, Axmacher, & Henson, 2012). These results argue in favor of a phase information coding mechanism along the long-axis and a multifaceted, non-uniform engagement along the long-axis for novelty processing. However, there remains a possibility that these discrete changes could be due to the electrodes being located in different layers of the hippocampus at this anterior to medial transition. Given that linear electrodes are placed within a structure that curves in the medial-lateral and rostral-caudal directions, adjacent contacts where the abrupt changes were detected could be located in different layers leading to inverted polarities and observed phase offsets.

The HH-HB locus is relevant from an anatomical perspective: The two major longitudinal association fiber systems in the hippocampal formation show extensive axon divergence within the dorsal two-thirds and within the ventral third of rat hippocampus, but few fibers cross between these subdivisions in rats (Amaral & Witter, 1989; Fricke & Cowan, 1978; Ishizuka et al., 1990; Li et al., 1994; Swanson et al., 1978). In monkeys, there are extensive interconnections in the posterior two-thirds of the hippocampus, which are more limited in its anterior one-third, although the boundary, in terms of intrinsic connectivity, is less clearly demarcated than in rodents (Kondo et al., 2009). It will be important to determine whether intrinsic anatomically connectivity in humans shows a similar partition as described above, and whether the decrease in coherence coincides with its anatomical locus on the long axis.

## Conclusion

On the basis of previous human fMRI and PET data (Grady, 2020), we hypothesized greater anterior *vs*. posterior hippocampal responses to unexpected stimuli. Our results, however, demonstrate that unexpected stimuli engage not only the anterior hippocampus (HH), but also middle (HB) and posterior (HT) portions. There is therefore no clear electrophysiological explanation in terms of iERP amplitude or oscillatory response for the anterior hippocampal response to novelty commonly observed with functional neuroimaging, although a local generator of the iERP to unexpected stimuli between hippocampal head and body (in the anterior portion of the hippocampus) was indicated by an iERP polarity inversion. Note that there are several types of novelty, including violation of expectation (i.e., prediction error), second order effects (initial experience of expectation violation), and contextual novelty (recency of prior occurrence). Whereas the current study focused on the first type, human fMRI studies show that anterior hippocampal responses habituate to repeated occurrences of unexpected stimuli (*i.e.,* second order effects; (B. A. Strange & Dolan, 2001; Yamaguchi, Hale, D’Esposito, & Knight, 2004), raising the possibility that a hippocampal head-specific electrophysiological response to unexpected stimuli would only be observed for the initial unexpected trials. By contrast, higher theta power to unexpected stimuli in HT could be relevant to studies implicating the posterior hippocampus in novelty responses (Kirchhoff, Wagner, Maril, & Stern, 2000; Knight, 1996; Rombouts et al., 1997; Stern et al., 1996).

Moreover, our observations provide new insights for a proposed model of functional organization along the human hippocampal long-axis of different superimposed patterns – gradual changes and discrete transitions (reviewed in Strange et al., 2014) – by demonstrating gradual increases in theta power from anterior to posterior, and discrete changes in neurophysiological properties at the junction between the HH and HB (**Figure 8**).

**Figure 8.**
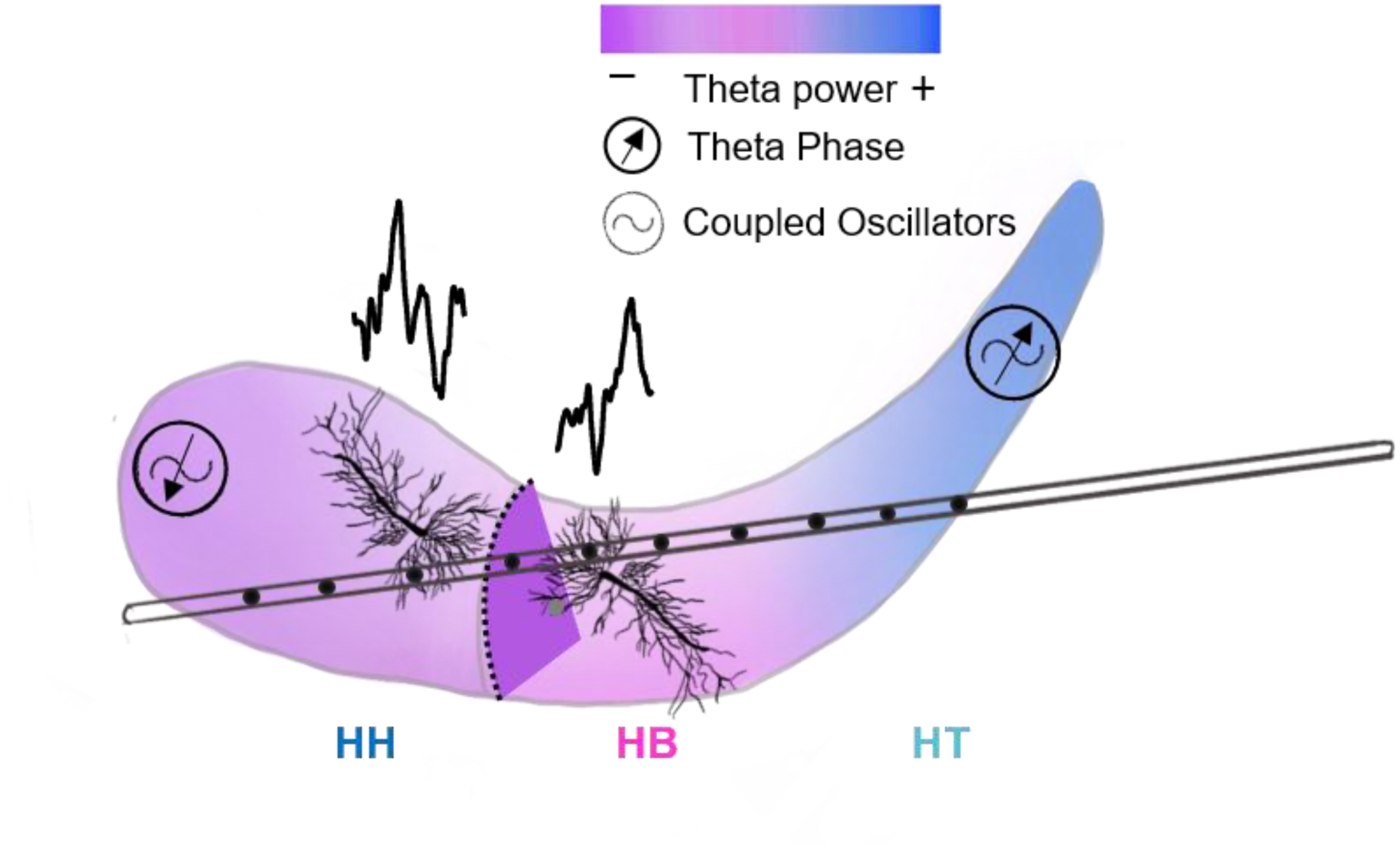
Summary schematic of hippocampal long-axis findings. Transition between HH and HB is indicated by dotted line. Though defined arbitrarily as the uncal apex, this transition coincides with a pronounced change of phase angle in theta and alpha traveling waves (indicated by the arrow within circle), as well as a polarity inversion in the evoked iERP to unexpected stimuli (example patient average traces either side of the HH-HB transition are shown above the hippocampus). The possibility that these effects reflect a change in layers, with different neuronal orientation, is illustrated by neurons at opposing orientations at the HH-HB transition.

## Author contributions

N.A. designed the experiment and collected data. M.Y. performed data analysis with guidance from O.J., N.A., S.M. and B.A.S. L.K provided electrode localization and imaging data. M.Y. and B.A.S. wrote the paper with input from all of the other authors.

## Data and materials availability

All data needed to evaluate the conclusions in the paper are present in the paper and/or the supplementary materials. The data and the code used to analyze data during the current study are available from the corresponding author on reasonable request.

## Supplementary Materials

**Supplementary Figure 1.**
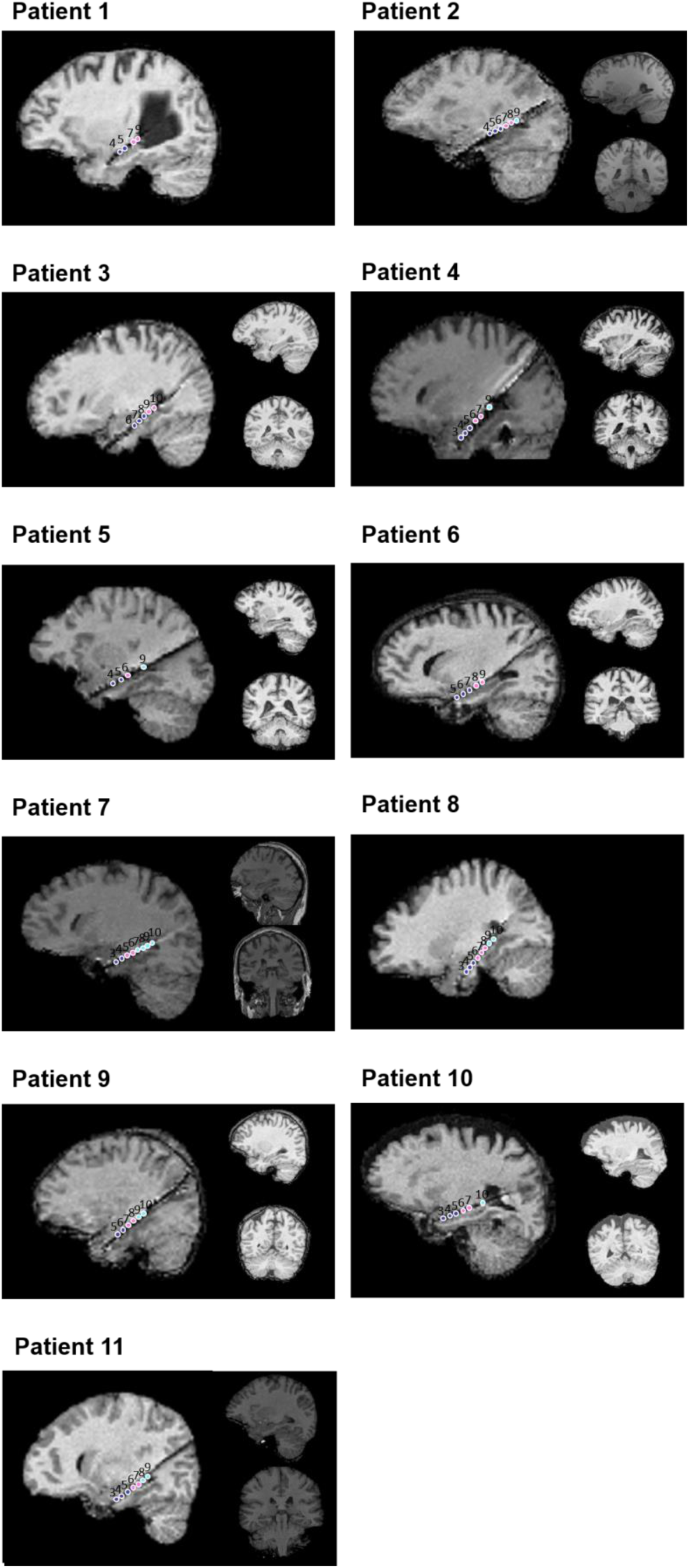
Pre- and post-implantation MRI for each patient. Pre- (two right panels; where available) and post-implantation (left panel) MRI showing the location of electrode contacts along the long axis of the hippocampus for each patient. Contacts within different regions of the hippocampus are depicted in different colors: dark blue (HH), pink (HB) and light blue (HT). Note that the pre-operative MRI scans for patients 1 and 8 are not available.

**Supplementary Figure 2.**
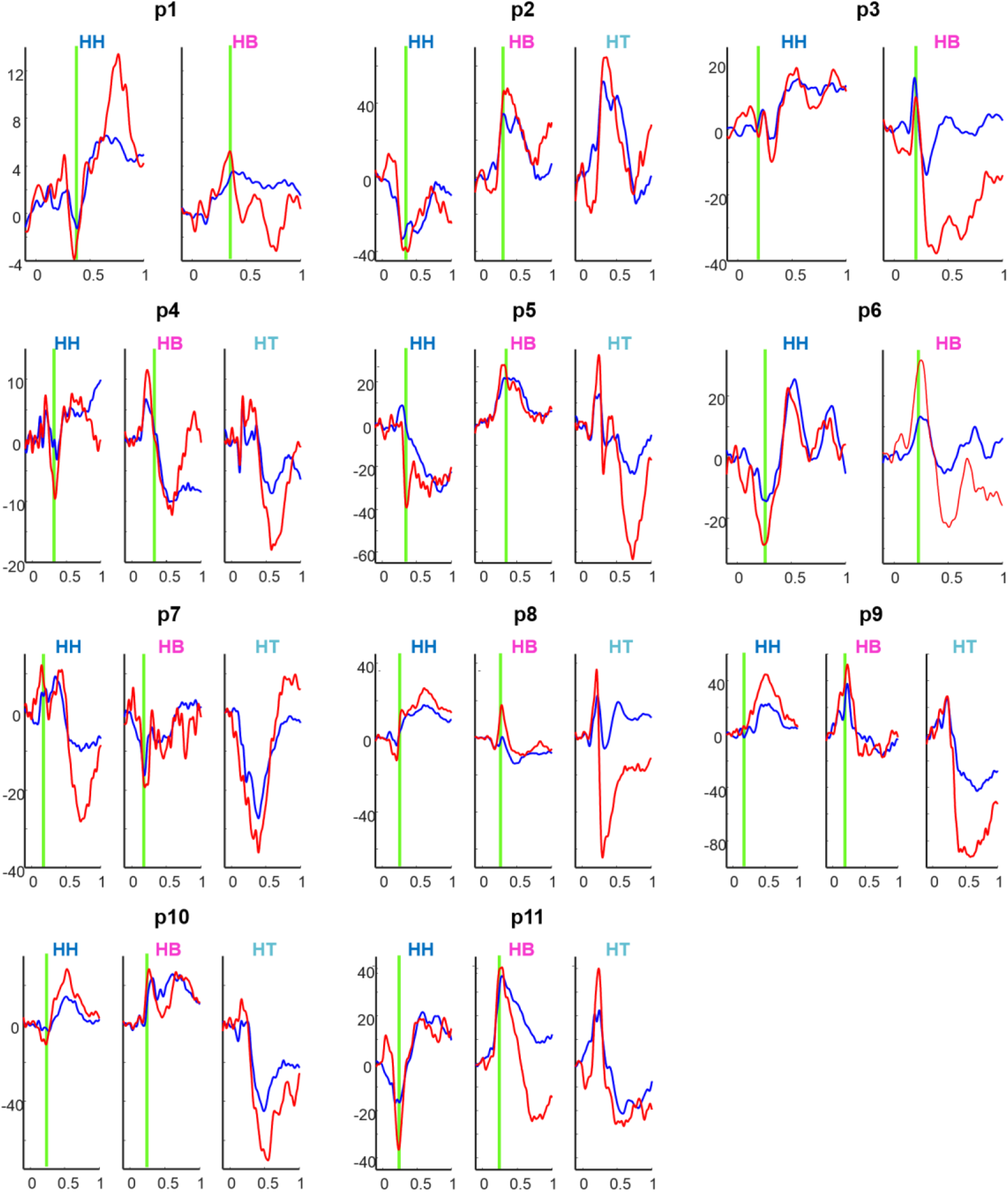
ERPs to unexpected (red) and expected (blue) stimuli for every subject and averaging contacts for each portion HH, HB and HT. The green vertical line indicates the polarity inversion.

**Supplementary Figure 3.**
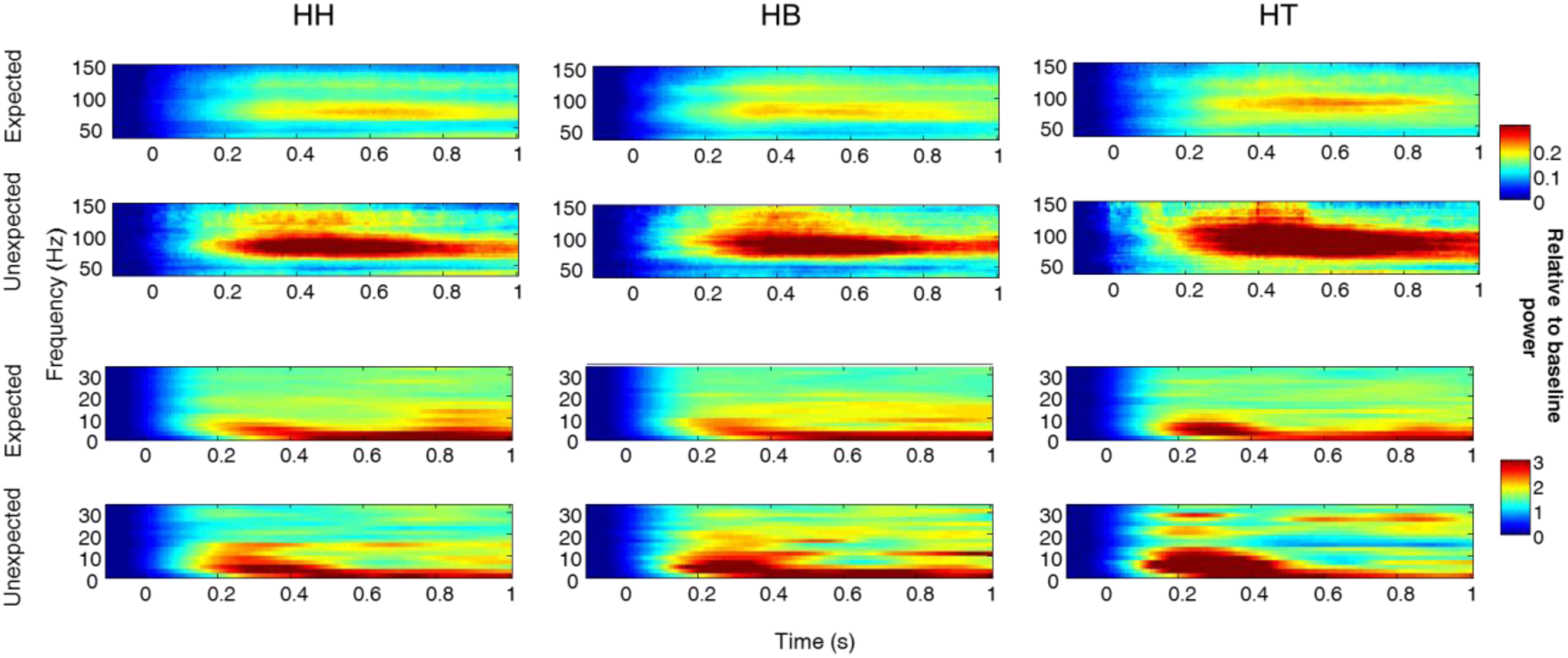
Time frequency plots for the relative to baseline power change for expected and unexpected conditions separately (baseline from -100ms to -10ms). Each portion HH, HB and HT in columns and for lower frequencies (0-30Hz) lower row and higher frequencies upper row (30-150Hz).

**Supplementary Figure 4.**
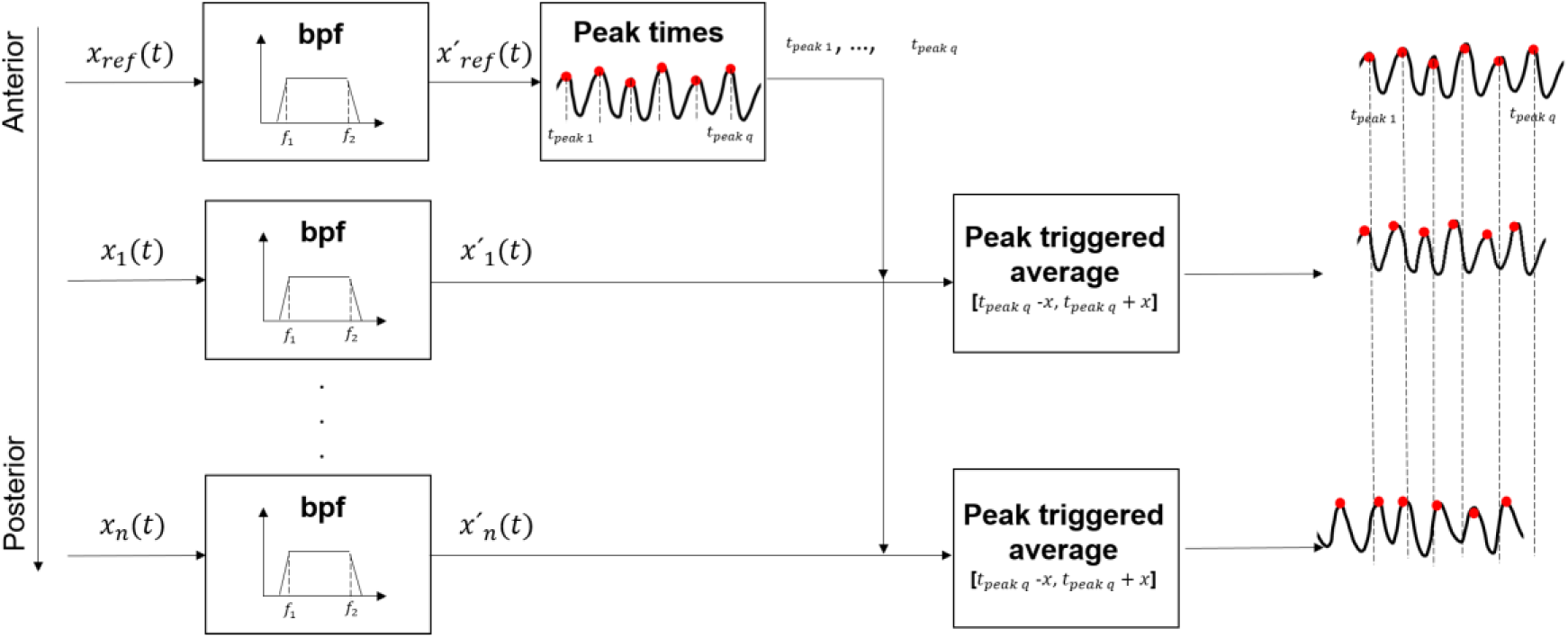
Travelling waves analysis approach 1. Signal from every hippocampal channel *x*_*n*_(*t*) was band pass filtered in the frequency bands of interest (theta or gamma) leading to *x*′_*n*_(*t*). In this approach, the filtered signal from the most anterior contact (first contact in the head) was considered the reference *x*_ref_(*t*). Time points at which the peaks in the filtered signal were identified (*t*_peak q_) and used as triggers to average the signal in the filtered signals from the rest of the hippocampal contacts. Data locked to the peaks of the reference channel (500 ms before and after the peaks of the reference channel) was averaged across trials for each channel. The resulting band pass filtered signal locked to the reference channel for each contact were plotted (Figure 6 and 7 C1 and C2) to compare peak times to the reference channel. If a traveling wave exists, time points at the peaks of the filtered signals, should increase or decrease consistently from the reference channel as the distance with it increases along the hippocampal longitudinal axis.

**Supplementary Figure 5.**
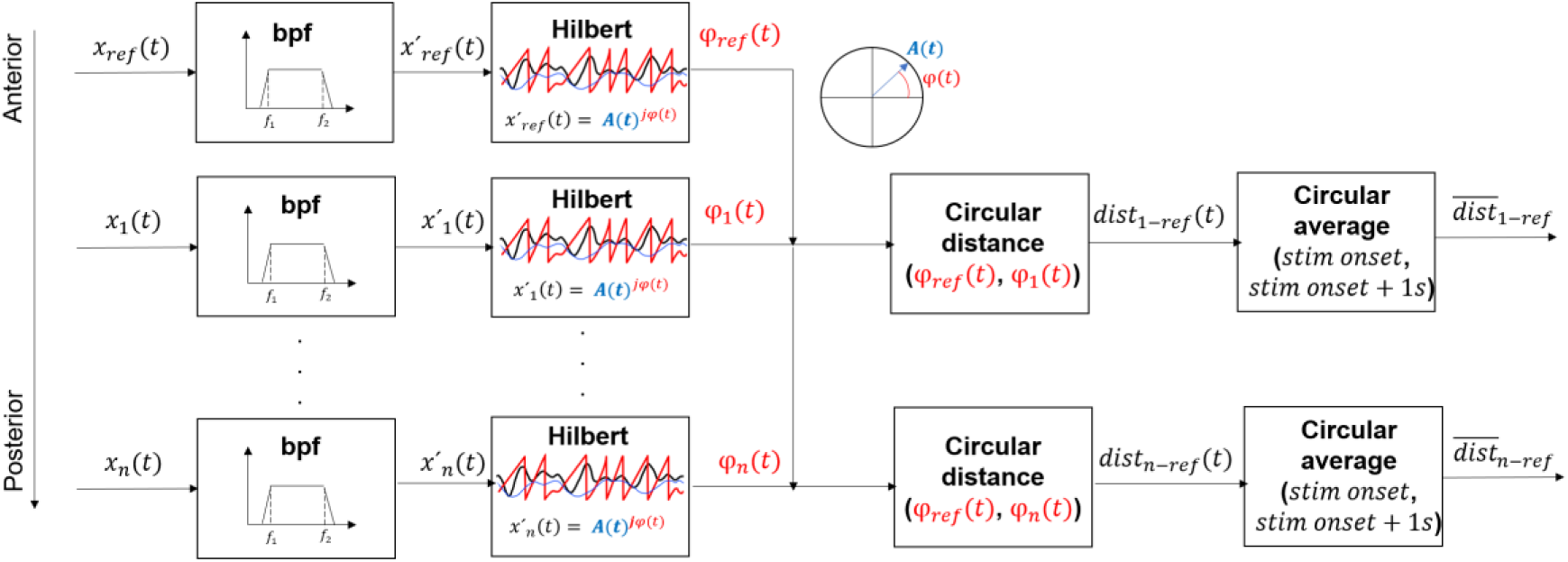
Travelling waves analysis approach 2. Signal from every hippocampal channel *x*_*n*_(*t*) was band pass filtered in the frequency bands of interest (theta or gamma) leading to *x*′_*n*_(*t*). In this approach, the filtered signal from the most anterior contact (first contact in the hipocampal head) was considered the reference *x_ref_*(*t*). Then, the analytic signal was calculated and the phase of the signal *φ*_*n*_(*t*) was computed as the angle from the Hilbert transform depicted in red. For every channel the angular distance with respect to the reference channel (the most anterior) *dist_ref−n_*(*t*) was computed using circular statistics (MATLAB toolbox CircStat2012a) and averaged across time 1s after stimulus onset for each trial to get *dist_ref−n_*. Polar plots depicted in Figures 6 and 7 D1 and D2, depict circular histograms of the computed phase distance in blue, and the phase locking value (plv; circular mean) across trials in red.

**Supplementary Figure 6.**
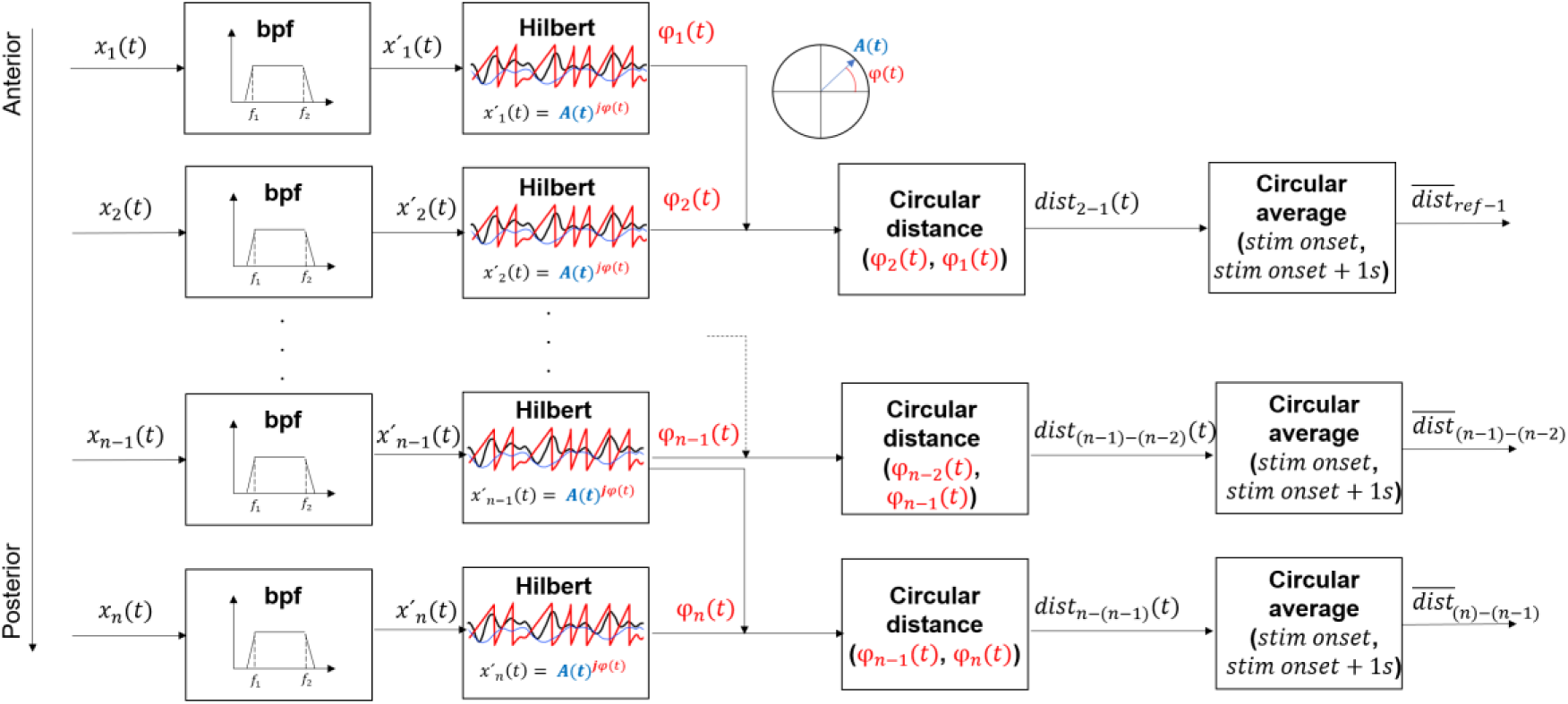
Travelling waves analysis approach 3. Signal from every hippocampal channel *x*_*n*_(*t*) was band pass filtered in the frequency bands of interest (theta or gamma) leading to *x*′_*n*_(*t*). In this approach, the filtered signal from the inmediate more anterior contact was considered the reference *x*_ref_(*t*). Then, the analytic signal was calculated and the phase of the signal *φ*_*n*_(*t*) was computed as the angle from the Hilbert transform depicted in red. For every channel the angular distance with the previous channel (more anterior) *dist*_*n*−(*n*−1)_(*t*) was computed using circular statistics (MATLAB toolbox CircStat2012a) and averaged across time 1s after stimulus onset for each trial to get *dist*_*n*−(*n*−1)_. Polar plots depicted in Figures 6 and 7 E1 and E2, depict circular histograms of the computed phase distance in blue, and the plv across trials in red.

**Supplementary Table 1.** Patient demographic and clinical data. F/M female/male; R right; - data unavailable

| Patient | Sex | Hand-<br>edness | Age<br>(years) | Age at<br>epilepsy<br>onset<br>(years) | Aetiology | Lesion location | Drugs dose (mg) | % of visually<br>rejected<br>trials |
| --- | --- | --- | --- | --- | --- | --- | --- | --- |
| 1 | M | R | 34 | - | Complex-focal and secondarily generalised seizures. Ammons Horn sclerosis right side. Medial temporal lobe epilepsy. Neurofibromatosis Type 1 with chiasma glioma. | Ammons Horn sclerosis right side. | Levetiracetam 1500-0-1500mg; Clobazam 0-0-5mg; L-Thyroxine 75-0-0µg | 0 |
| 2 | M | R | 28 | 10 | Simple- and complex-partial and secondarily generalised seizures. Suspected disorder of left medial temporal grey-white matter differentiation. Depressive disorder. | Anterior left temporal lobe. | Lamotrogine 200-0-200mg; Clobazam 0-0-5mg | 15 |
| 3 | F | R | 33 | - | Complex-partial seizures. Left Ammons Horn sclerosis. | Ammons Horn sclerosis left side (left hippocampus). | Levetiracetam 1500-0-2000mg; Clobazam 5-0-5mg; | 12 |
| 4 | F | R | 38 | - | Cryptogenic temporal lobe epilepsy with complex-partial seizures. | Left temporal (electrodes TL, TBAL, TBPL). | Levetiracetam 1500-0-1500mg; Clobazam 5-0-5mg; | 10 |
| 5 | F | R | 48 | 18 | Cryptogenic epilepsy with complex-partial seizures. | Particularly left temporo-polar. | Lamotrigine 200-0-200mg; Potassium Iodide 200-0-0mg; Clobazam 10-0-10mg | 11 |
| 6 | F | R | 45 | - | Cryptogenic focal epilepsy with complex-partial and secondarily generalised seizures with temporal semiology. | Temporal right. | Lamotrogine 50-0-50mg; Zonisamide 100-0-100mg; | 16 |
| 7 | M | R | 46 | 10 | Focal epilepsy with complex-partial seizures. Left sided Ammons Horn Sclerosis. Cortical disorder of differentiation left temporal pole within the gyrus temporalis inferior. | Ammons Horn sclerosis left side. | Carbamazepine ret. 400-0-800mg; Zonisamide 100-0-100mg; | 21 |
| 8 | F | R | 25 | 6 | Cryptogenic epilepsy with complex-partial and secondarily generalised seizures. MR tomographic suspicious lesion temporopolar left. Folate deficient. | Temporal left (TLL 5-7). | Carbamazepine ret. 400-0-600mg; Folate 5-0-0mg; Opipramol 50-50-50mg; | 18 |
| 9 | F | R | 18 | - | Focal epilepsy with complex-partial seizures. Left Ammons Horn sclerosis. Cognitive impairment. | Ammons Horn sclerosis left side. | Oxcarbazepine 900-0-900mg; Clobazam 5-0-5mg | 14 |
| 10 | F | R | 56 | 34 | Cryptogenic epilepsy with right temporomesial starting complex-partial seizures. | Right temporo-mesial. | Levetiracetam 1500-0-1500mg | 31 |
| 11 | F | R | 29 | - | Epilepsy with complex-partial seizures. Right Ammons Horn sclerosis and dysplasia of the right temporal pole. | Ammons Horn sclerosis right side. | Levetiracetam 1500-0-1500mg; Lamotrogine 150-0-150mg; Clobazam 10-0-10mg; | 25 |

**Supplementary Table 2.** Labels of the channels used for the absolute Coherence and Coherence angle analysis between portion.

|  | Between portions |  |  |
| --- | --- | --- | --- |
|  | HH-HB | HH-HT | HB-HT |
| **P 1** | HH1-HB1<br>HH1-HB2<br>HH2-HB1<br>HH2-HB2 | -- | -- |
| **P 2** | HH1_HB1<br>HH2_HB1<br>HH3_HB1<br>HH1_HB2<br>HH2_HB2<br>HH3_HB2 | HH1_HT<br>HH2_HT<br>HH3_HT | HB1_HT<br>HB2_HT |
| **P 3** | HH1_HB1<br>HH2_HB1<br>HH3_HB1<br>HH1_HB2<br>HH2_HB2<br>HH3_HB2 | -- | -- |
| **P 4** | HH1_HB1<br>HH2_HB1<br>HH3_HB1<br>HH1_HB2<br>HH2_HB2<br>HH3_HB2 | HH1_HT<br>HH2_HT<br>HH3_HT | HB1_HT<br>HB2_HT |
| **P 5** | HH2_HB<br>HH3_HB | HH2_HT<br>HH3_HT | HB_HT |
| **P 6** | HH1_HB1<br>HH2_HB1<br>HH3_HB1<br>HH1_HB2<br>HH2_HB2<br>HH3_HB2 | -- | -- |
| **P 7** | HH1_HB1<br>HH2_HB1<br>HH1_HB2<br>HH2_HB2 | HH1_HT1<br>HH2_HT1<br>HH1_HT2<br>HH2_HT2<br>HH1_HT3<br>HH2_HT3<br>HH1_HT4<br>HH2_HT4 | HB1_HT1<br>HB2_HT1<br>HB1_HT2<br>HB2_HT2<br>HB1_HT3<br>HB2_HT3<br>HB1_HT4<br>HB2_HT4 |
| **P 8** | HH1_HB1<br>HH2_HB1<br>HH3_HB1<br>HH1_HB2<br>HH2_HB2<br>HH3_HB2<br>HH1_HB3<br>HH2_HB3<br>HH3_HB3 | HH1_HT1<br>HH2_HT1<br>HH3_HT1<br>HH1_HT2<br>HH2_HT2<br>HH3_HT2 | HB1_HT1<br>HB2_HT1<br>HB3_HT1<br>HB1_HT2<br>HB2_HT2<br>HB3_HT2 |
| **P 9** | HH1_HB1<br>HH2_HB1<br>HH1_HB2<br>HH2_HB2 | HH1_HT1<br>HH2_HT1<br>HH1_HT2<br>HH2_HT2 | HB1_HT1<br>HB2_HT1<br>HB1_HT2<br>HB2_HT2 |
| **P 10** | HH1_HB<br>HH2_HB<br>HH3_HB<br>HH4_HB | HH1_HT<br>HH2_HT<br>HH3_HT<br>HH4_HT | HB_HT |
| **P 11** | HH1_HB1<br>HH2_HB1<br>HH3_HB1<br>HH1_HB2<br>HH2_HB2<br>HH3_HB2 | HH1_HT1<br>HH2_HT1<br>HH3_HT1<br>HH1_HT2<br>HH2_HT2<br>HH3_HT2 | HB1_HT1<br>HB2_HT1<br>HB1_HT2<br>HB2_HT2 |

**Supplementary Table 3.** Summary for the cluster-based statistical analysis between expected and unexpected conditions per each portion. Minimum and maximum frequencies and time windows for the significant clusters. p_clust_ p values corrected for multiple comparisons.

| Freq range | Cluster results | Portion |  |  |
| --- | --- | --- | --- | --- |
|  |  | HH | HB | HT |
| **Low**<br>2-34 Hz | [tmin,tmax] (ms) | [160 420] | [180 380] | [0 450] |
|  | [fmin, fmax] (Hz) | [3 5] | [3 5] | [1 11] |
| | $p_{clust}$ | 0.025 | 0.012 | 0.0009 |
| **High**<br>34-150 Hz | [tmin,tmax] (ms) | [120 750] | [60 800] | [90 1000] |
|  | [fmin, fmax] (Hz) | [60 90] | [67 112] | [80 150] |
| | $p_{clust}$ | 0.003 | 0.0009 | 0.0009 |

**Supplementary Table 4.**
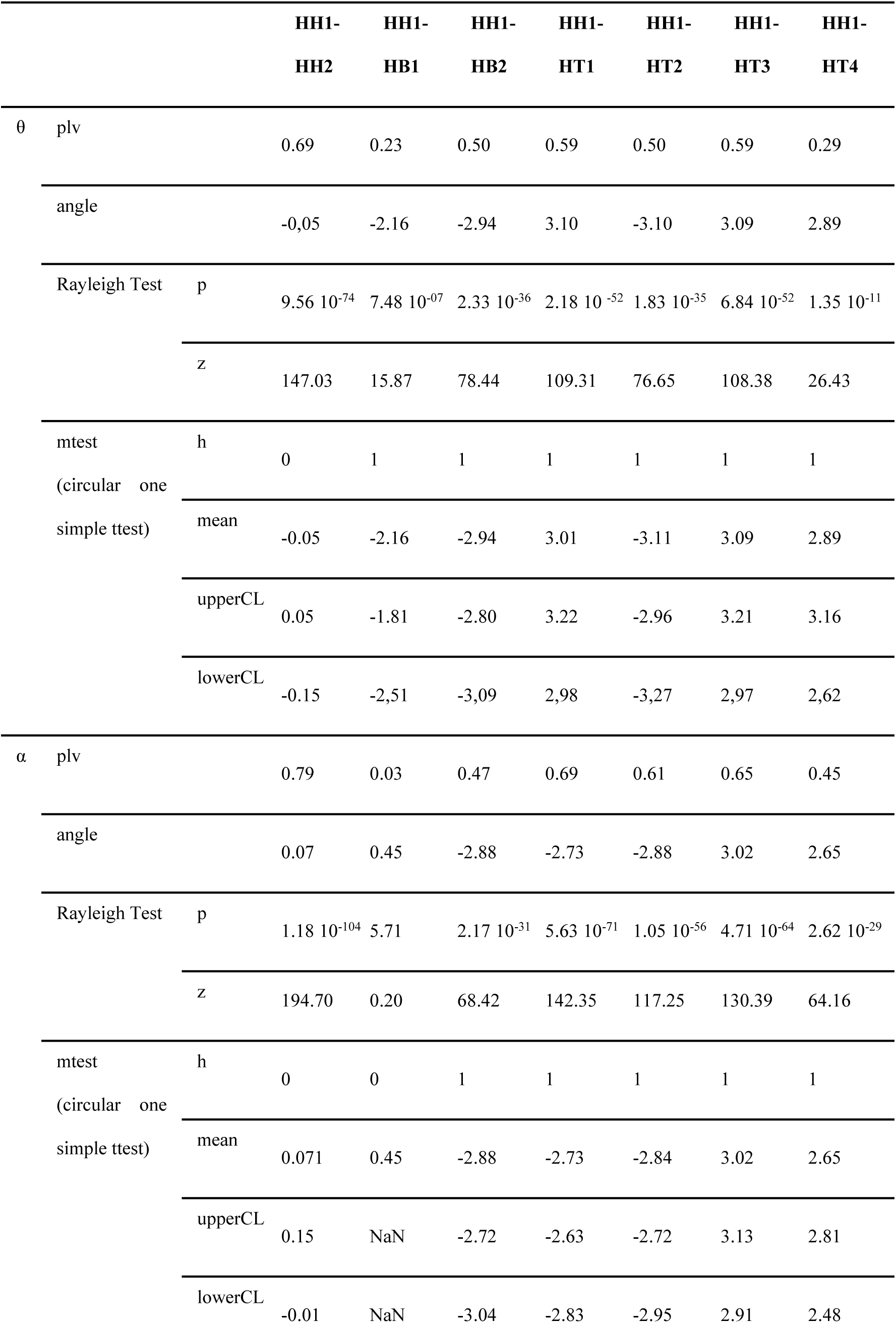
Statistics for the phase distance of every contact with the most anterior as a reference for patient 7 (phase angle in radians). h: null hypothesis, h=0 indicates that h cannot be rejected at the 5% significance level, h=1 that it can be rejected. CL: confidence limits

**Supplementary Table 5.**
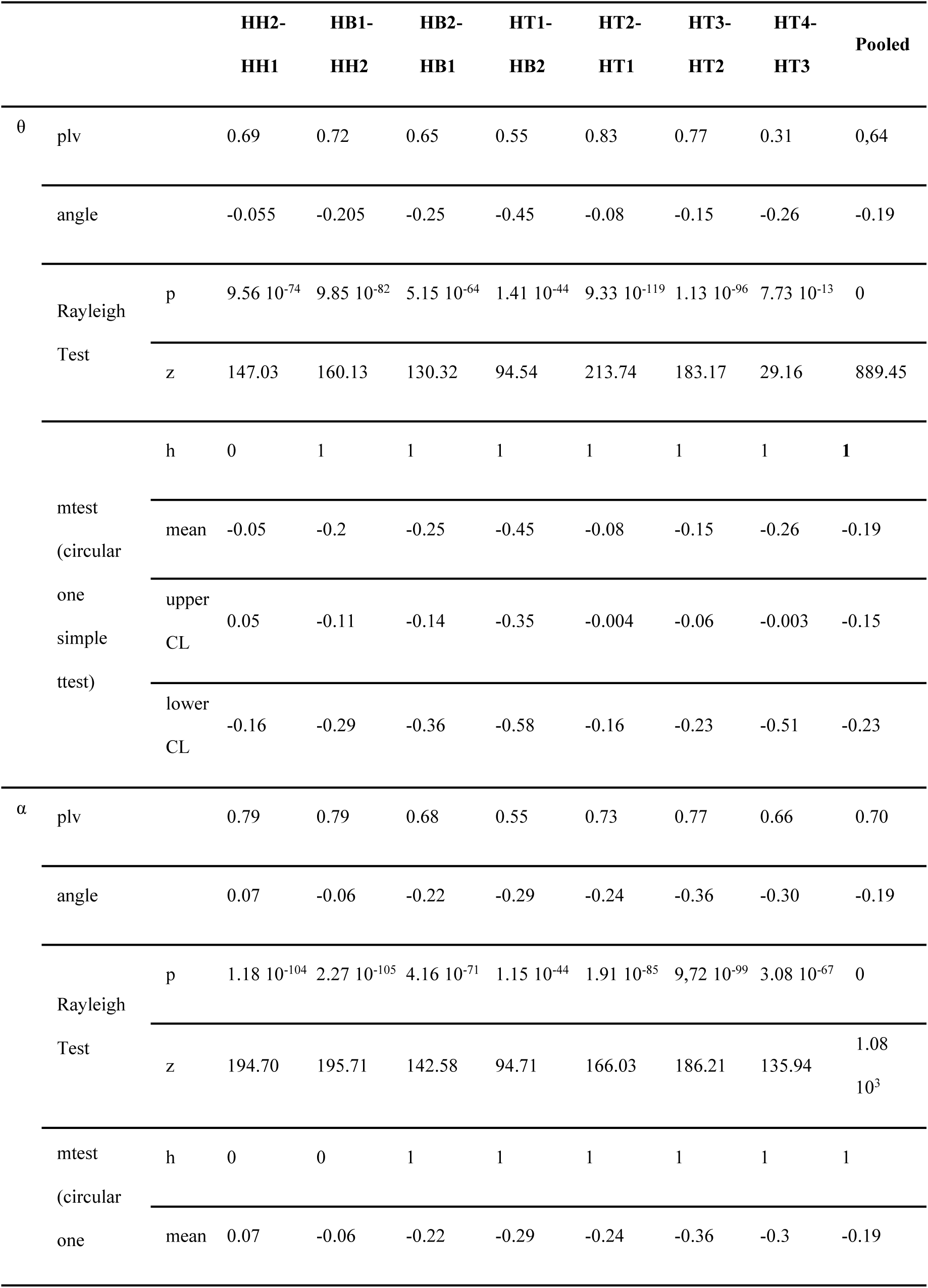
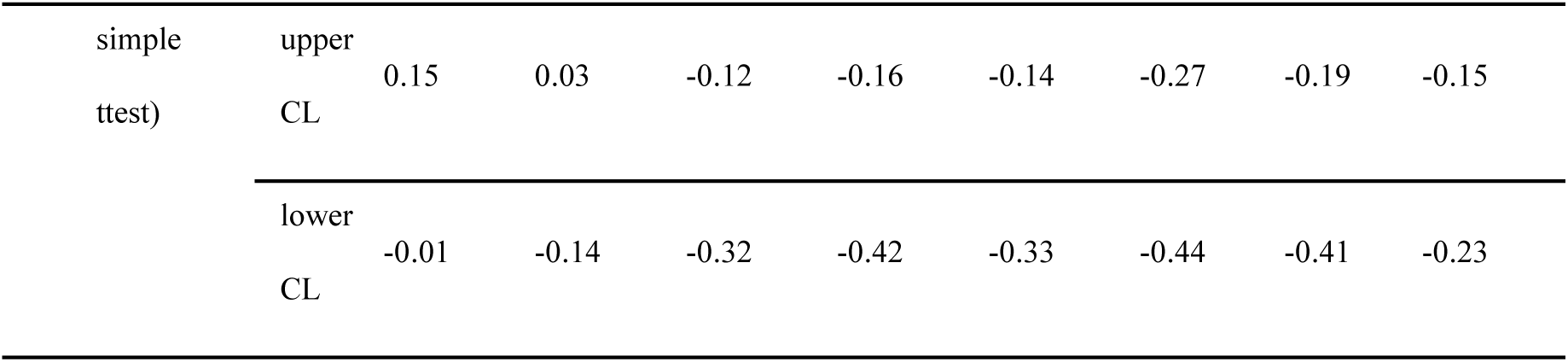
Statistics for the phase distance of every pair of adjacent contact for patient 7 (phase angle in radians). h: null hypothesis, h=0 indicates that h cannot be rejected at the 5% significance level, h=1 that it can. CL: confidence limits

**Supplementary Table 6.**
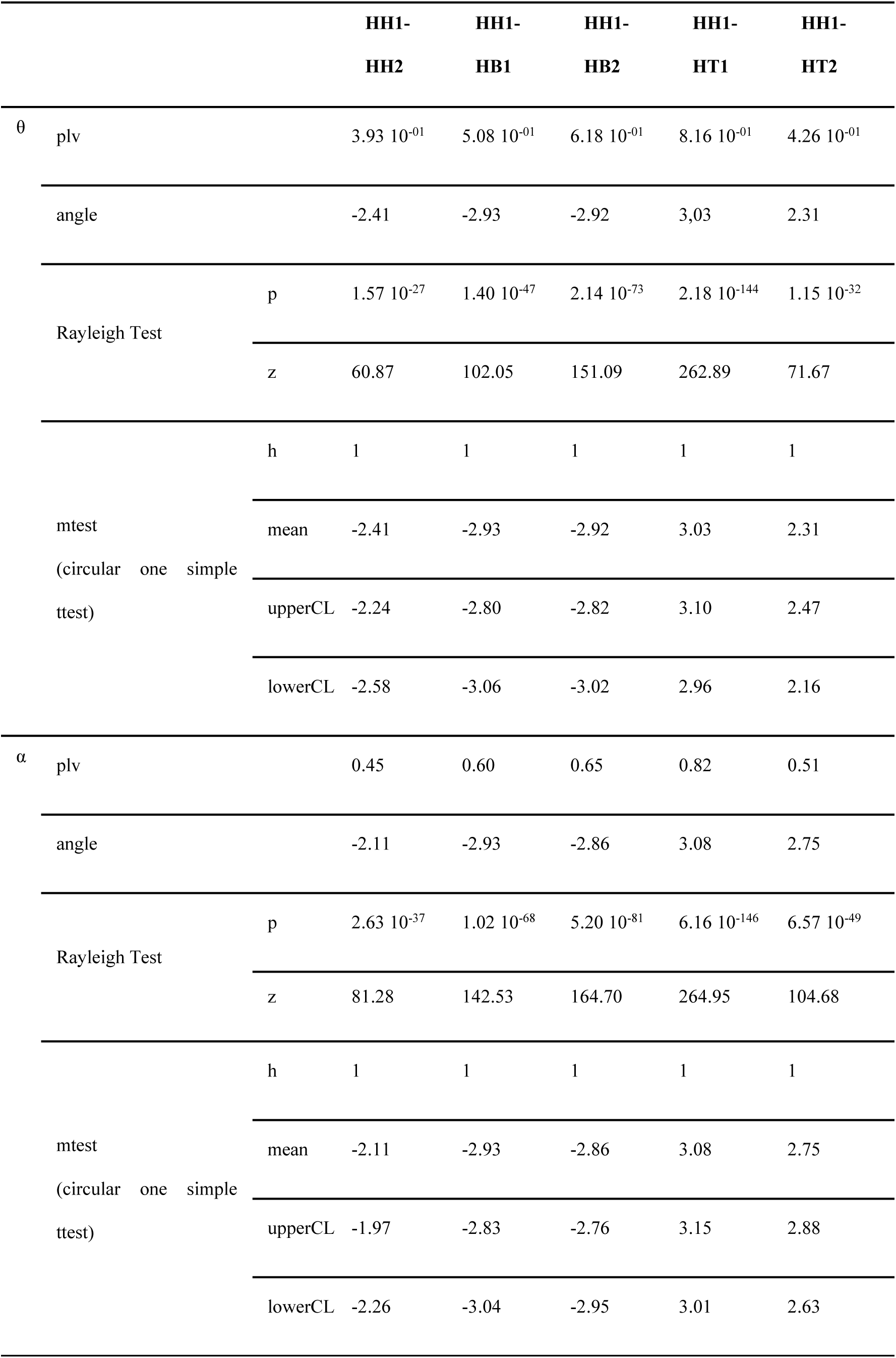
**Supplementary Table 6**. Statistics for the distance of every contact with the most anterior as a reference for patient 9. (phase angle in radians). h: null hypothesis, h=0 indicates that h cannot be rejected at the 5% significance level, h=1 that it can. CL: confidence limits

**Supplementary Table 7.**
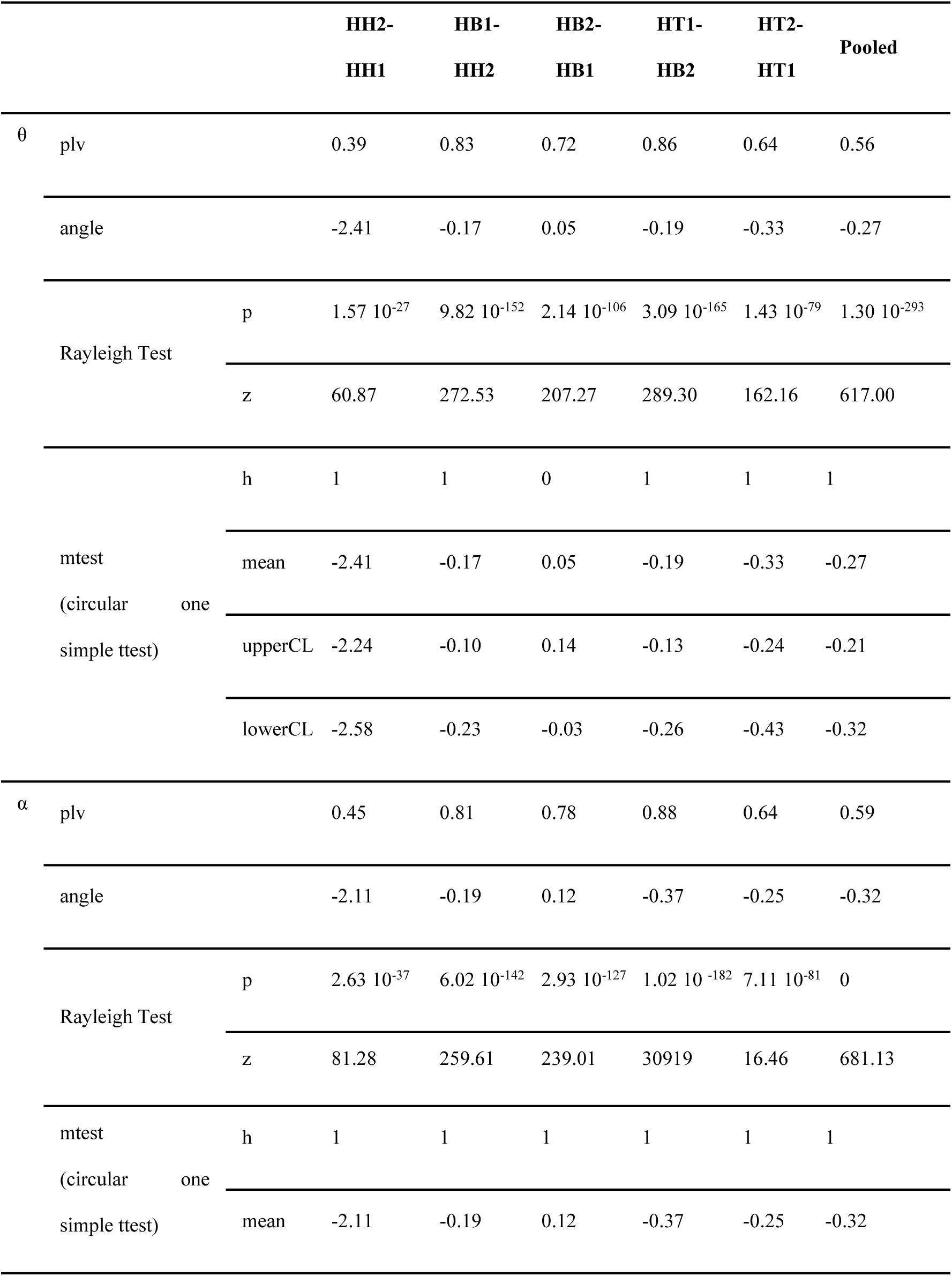
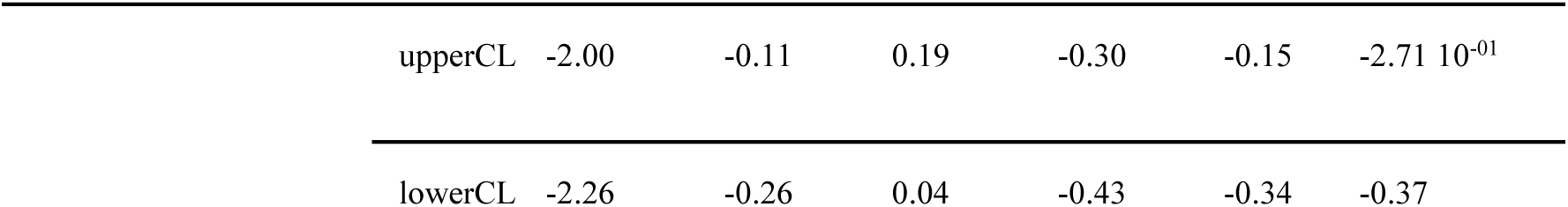
Statistics for the distance of every pair of adjacent contact for patient 9 (phase angle in radians). h: null hypothesis, h=0 indicates that h cannot be rejected at the 5% significance level, h=1 that it can. CL: confidence limits

### Supplementary Results. Hippocampal head to body transition is not associated with consistent subfield transition

The implantation of depth electrodes along the hippocampal long-axis can lead to variability in exact medio-lateral and infero-superior position of each electrode contact within and between hippocampal portions. This raises the possibility that transitions between portions coincide with transitions between hippocampal subfields. If this were to occur consistently for electrode contacts on either side of the HB to HH border as defined here, the polarity inversion of novelty evoked iERPs, the discrete change in coherence and the sharp phase transition in long-axis traveling waves could, potentially, be attributed to a subfield transition. Indeed, simultaneous recordings at multiple depths in rat dorsal hippocampus have shown an abrupt drop in theta coherence occurs across subfield borders (CA1-subiculum) as well as less extreme drops within subfields (Bullock, Buzsaki, & McClune, 1990). Furthermore, phase profiles across CA1 and into subiculum show abrupt, local shifts of phase (Bullock et al., 1990). To control for this possibility, we ascribed electrode contact locations to specific subfields using a probabilistic atlas (Eickhoff et al., 2005). Across patients, there were 3 in whom HB-HH transitions comprised contacts in the same subfield (2 DG-DG and 1 CA1-CA1), three with transitions from CA3 to CA1, one with transitions from CA2 to CA1, one hippocampus to CA3 and 2 for whom probabilistic localization failed for one or both contacts. Thus, although the approach of assigning anatomy with a probabilistic atlas is not without its caveats (Eickhoff et al., 2007), we do not find a consistent subfield change at the anatomical locus where we observe a discrete change in coherence and a sharp phase transition in long-axis traveling waves, suggesting that these changes are more likely to arise from a change in some property of theta (and alpha) oscillations along the long-axis. There remains, nonetheless, the possibility that aspects of our findings relate to subfield-specific phase shifts.

